# Mapping the gene regulatory landscape of archaic hominin introgression in modern Papuans

**DOI:** 10.1101/2025.05.04.652069

**Authors:** Maddy Comerford, Davide M. Vespasiani, Navya Shukla, Laura E. Cook, Danat Yermakovich, Michael Dannemann, Matthew Leavesley, Christopher Kinipi, François-Xavier Ricaut, Nicolas Brucato, Murray P. Cox, Irene Gallego Romero

## Abstract

Interbreeding between anatomically modern humans and archaic hominins has contributed to the genomes of present-day human populations. However, our understanding of the specific gene regulatory consequences of Neanderthal, and particularly, Denisovan introgression is limited. Here, we used a massively parallel reporter assay to investigate the regulatory effects of 25,869 high-confidence introgressed SNPs segregating in present-day individuals of Papuan genetic ancestry in immune cell types. Overall, 8.22% of Denisovan and 8.58% of Neanderthal sequences showed active regulatory activity, and 9.22% of these displayed differential activity between alleles. We examined the properties of active and differentially active sequences, and found no impact of introgressed allele frequency on the probability of activity, but do observe an association with distance to the nearest transcription start site. Differential activity was also associated with differential transcription factor binding. Genes predicted to be regulated by differentially active sequences included *IFIH1* and *TNFAIP3*, key immune genes and known examples of archaic introgression. Overall, this work provides insight into the regulatory activity of archaic variants in Papuan populations and, more broadly, the contribution of archaic introgression in shaping modern human genetic diversity.

## Introduction

As modern humans left Africa and spread across the globe, they came into contact with multiple archaic human groups, two of which we have published genome sequences for: Denisovans (Reich et al., 2010; Meyer et al., 2012) and Neanderthals (Green et al., 2008; Prüfer, Racimo, et al., 2014; Prüfer, de Filippo Cesare, et al., 2017; Mafessoni et al., 2020). Multiple interbreeding events between expanding anatomically modern humans (AMH) and archaic groups led to present-day humans carrying introgressed archaic DNA. Reflecting a complex population history, this introgression is not uniformly distributed across AMH populations today. Neanderthal introgression has contributed up to 2% to the genomes of individuals of non-African genetic ancestry (Prüfer, Racimo, et al., 2014), while Denisovan introgression accounts for up to 5% of the genomes of individuals of Papuan or Near Oceanic genetic ancestry (Reich et al., 2010; Meyer et al., 2012) and under 1% in individuals of South or East Asian genetic ancestry, and is virtually absent elsewhere in the world.

Additionally, archaic introgression is not distributed uniformly across the genome. While selection against introgressed DNA appears to have been widespread across the genome (Telis et al., 2020; Sankararaman et al., 2016), we and others have previously shown a significant excess of introgressed DNA at or near genes involved in immune function (Vespasiani et al., 2022; Quach et al., 2016; Dannemann, Andrés, et al., 2016; Nédélec et al., 2016; Gittelman et al., 2016; Deschamps et al., 2016; Sams et al., 2016; Villanea et al., 2023; Abi-Rached et al., 2011; Mendez et al., 2012; Mendez et al., 2013; André et al., 2024; Yermakovich et al., 2024), suggesting that introgression was not always disadvantageous. Particularly in the case of immune phenotypes, introgressed alleles would have been ’fine tuned’ relative to those carried by expanding AMH, and better suited to countering the threats posed by local pathogens. Their retention would thus have conferred a selective advantage to carriers (Abi-Rached et al., 2011). Much of this advantage is thought to be mediated not by changes to protein coding genes (although there are notable exceptions, e.g. Zammit et al., 2019) but primarily by driving fine-scale changes in gene expression levels. For example, Neanderthal sequences are overrepresented amongst steady-state and response eQTLs (Dannemann, Prüfer, et al., 2017; Quach et al., 2016). Additionally, introgressed Neanderthal sequences have been predicted to impact allele-specific gene expression levels (McCoy et al., 2017) and to affect enhancers in a tissue-specific manner (Silvert et al., 2019).

However, our understanding of the specific functional consequences of Denisovan introgression in present-day human populations remains limited. In comparison to Neanderthal introgression, Denisovan introgression is under-represented in genomic databases, because Denisovan DNA is not present in the genomes of European populations who make up the large majority of genomic databases (Mills and Rahal, 2020), and much introgression falls in non-coding parts of the genome where function is harder to predict directly from sequence. While we previously showed that in the genomes of present-day Papuan individuals, both Neanderthal and Denisovan introgressed alleles are enriched within potential tissue-specific gene regulatory elements, and in particular within chromatin regions active in various immune cell types (Vespasiani et al., 2022), we could not validate these predictions against actual gene expression data due to the near complete absence of individuals of Papuan genetic ancestry from gene expression datasets (Natri et al., 2022).

Massively parallel reporter assays (MPRAs) allow for functional testing of regulatory activity at scale, and are rapidly becoming part of the evolutionary and comparative genetics toolkit (Gallego Romero and Lea, 2023). In an MPRA, a synthesised library of short synthetic oligos predicted to regulate gene expression levels is tagged with unique barcodes and cloned in front of a minimal promoter and a reporter gene, such that the functional elements will drive transcription of the barcode and reporter gene (Melnikov, Murugan, et al., 2012). In this manner, thousands, hundreds of thousands, or millions of sequences can be tested simultaneously, overcoming the low throughput of traditional reporter assays (Inoue and Ahituv, 2015). While initially applied to biomedical questions, MPRAs are increasingly being used to understand evolutionary questions (e.g., Jagoda et al., 2022; Uebbing et al., 2021; Weiss et al., 2021; Findley et al., 2021). Particularly in light of the systematic under-representation of peoples of non-European genetic ancestry in public genetic and cellular resources, MPRAs provide a tractable and powerful way to examine the gene regulatory impact of archaic introgression in present-day Papuans.

Thus, to characterise the regulatory activity of archaic variants present in these individuals in immune cells, we have tested the activity of over 25,000 Denisovan and Neanderthal variants segregating in present-day Papuan populations at high frequencies. We identify Denisovan and Neanderthal alleles that robustly drive reporter gene expression, and by testing both the archaic and non-archaic allele of each variant, identify variants for which modern and introgressed alleles drive significantly different reporter gene expression. This work provides, for the first time, experimental insight into the regulatory activity of archaic variants segregating within Papuan populations and, more broadly, insights into the contribution of archaic introgression in shaping modern human genetic diversity.

## Results

### Identifying cis regulatory activity of introgressed sequences using an MPRA

To characterise the functional impact of archaic introgression on the gene regulatory landscape of modern-day Papuan populations, we tested the regulatory activity of 25,869 introgressed alleles and their modern human counterparts using an MPRA (Figure 1A). Starting with the set of introgressed SNPs identified in Papuan populations by Jacobs et al., 2019, we followed the same approach as in Vespasiani et al., 2022 and focused on those located in open chromatin, defined by DNAseI hypersensitive sites from (Kundaje et al., 2015; Thurman et al., 2012) or Tn5 transposase accesible regions from (Calderon et al., 2019), or in active chromatin states defined by the Roadmap Epigenome Project for at least one immune cell type (defined as all cells within HSC and B cells or blood and T cells in Kundaje et al., 2015; Supplementary Figures 1 and 2). We also included introgressed SNPs predicted to disrupt transcription factor binding sites (Vespasiani et al., 2022) (Figure 1B). Finally, we limited ourselves to variants with an introgressed allele frequency (IAF) ≥ 0.15 (Supplementary Figure 3), reasoning they were more likely to confer a selective advantage than those at lower frequencies. To quantify the activity of these variants we designed 200bp oligos containing 15bp flanking adapter sequences surrounding 170bp centred on the SNP of interest. For each allele, we included in our library an oligo containing the modern human allele, and an oligo containing the introgressed archaic allele. Where two or more SNPs were located within 170bp of each other, which happened 15.39% of the time, we additionally designed haplotype-like oligos containing all possible allelic combinations of the SNPs. In total, our initial MPRA library contained 61,812 sequences across 25,869 high-confidence introgressed SNPs. This included 35,198 and 25,496 single and haplotype-like sequences testing 14,775 and 11,094 Denisovan and Neanderthal SNPs, respectively, which below we refer to as Denisovan and Neanderthal sequences. We also included 153 SNPs previously found to be active in a previous MPRA (Tewhey et al., 2016) and 196 SNPs active in HepG2 cells (Arensbergen et al., 2019) to act as positive controls, and a set of 300 random scrambled sequences to act as negative controls. All tested sequences are available as **Supplementary file 1**.

**Figure 1.**
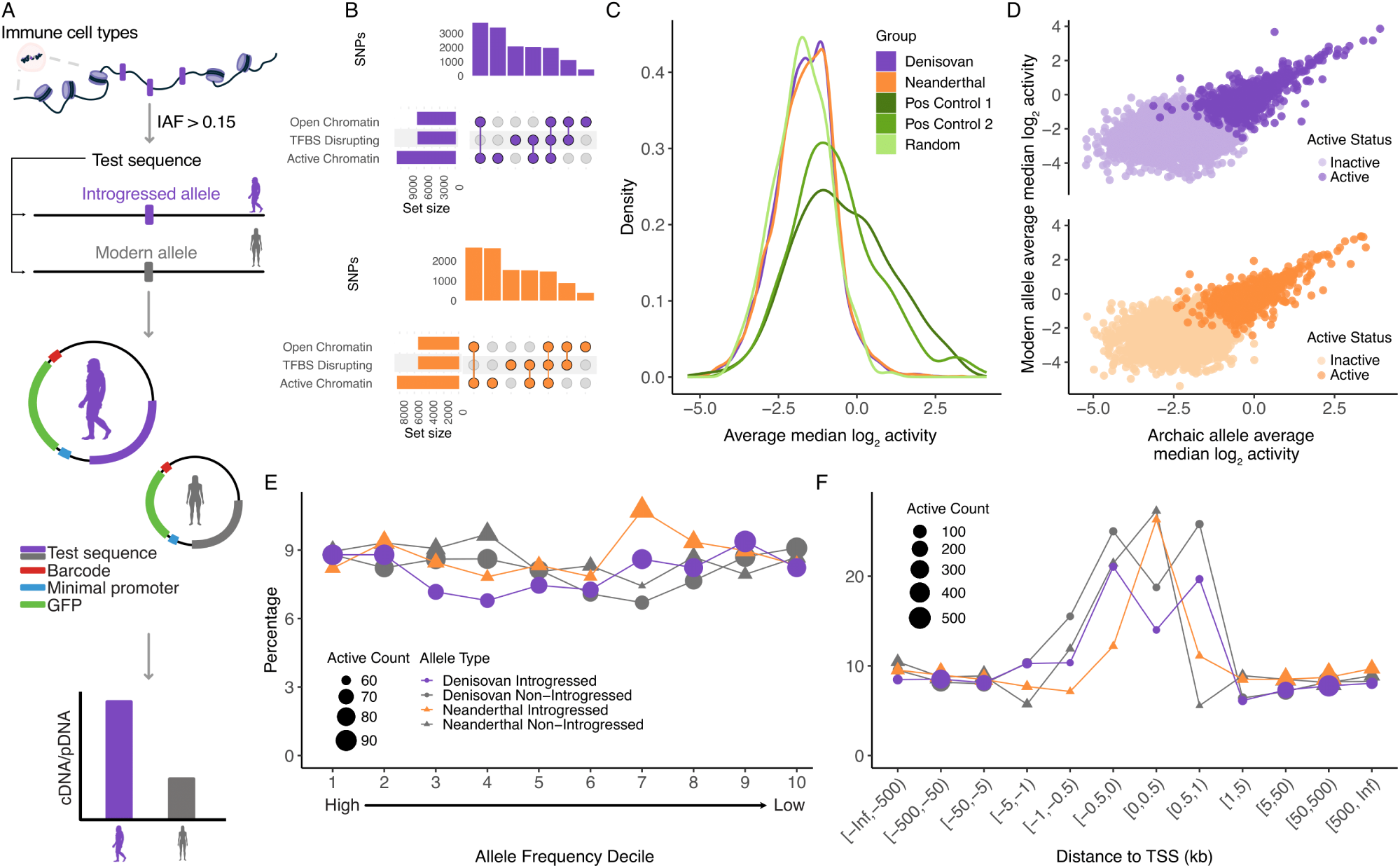
Quantifying the regulatory activity of archaic SNPs with an MPRA. A.) SNP selection and experimental overview. B.) Number of selected Denisovan (top) and Neanderthal (bottom) SNPs annotated across various functional annotations. C.) Distribution of activity for Denisovan, Neanderthal and control sequences. Positive control 1 and 2 sequences were previously found to be active by Tewhey et al., 2016 and Arensbergen et al., 2019, respectively. D.) Scatter plot of activity of the archaic allele and the modern allele for Denisovan (top) and Neanderthal (bottom) SNPs, with SNPs with at least one active allele highlighted. E.) Percentage of active alleles in allele frequency deciles. F.) Percentage of active alleles in binned distances to the transcription start site (TSS).

After barcode addition and initial library quantification (see Methods), we transfected the library into two replicates each of three different lymphoblastoid cell lines (GM12878, a European LCL and PNG8 and PNG22, established from Papuan donors sampled in Port Moresby). We collected both RNA and plasmid DNA (pDNA), and sequenced barcode counts in the cDNA and pDNA fractions, to an average depth of 247 million reads per library. We were unable to sequence cDNA from PNG8 replicate 1 due to low RNA recovery, so this replicate was discarded from downstream analyses. After filtering (see Methods), across replicates we captured a total of 44,144 sequences for activity testing, out of 59,954 (73.63%) of the sequences captured after library quantification. Measured sequences were associated with 48 barcodes on average (SD = 40, range = 10 to 562; Supplementary Figure 4). Sequence counts were moderately correlated between the cDNA and pDNA fractions within replicates (Spearman’s rank correlation *ρ* = 0.62 to 0.74, p <2.2 × 10^−16^), with a subset of sequences showing higher cDNA than pDNA counts (Supplementary Figure 5).

In total, 3,894 sequences (8.82% of measured sequences) showed consistent signs of regulatory activity in at least two replicates, defined as having a ratio of cDNA/pDNA counts across all barcodes that was significantly higher (FDR < 0.05) than the mean value for that replicate (see Methods). Positive control sequences showed higher activity than our archaic sequences and random scrambled sequences (Figure 1C), with 91 control sequences previously identified to be active in LCLs (Tewhey et al., 2016), and 154 control sequences previously identified to be active in HepG2 cells (Arensbergen et al., 2019) labelled as active (47.89% and 38.4%, respectively, of sequences tested in these groups), validating our experimental approach. Considering modern and introgressed alleles together, at least one allele in 2,252 tested sequences was active (**Supplementary file 2**); this includes 2,059 alleles across 1,254 Denisovan sequences and 1,568 alleles across 998 Neanderthal sequences. Both alleles were deemed active in 571 of tested Denisovan sequences and 457 of Neanderthal ones. Overall, 8.22% and 8.58% of tested Denisovan and Neanderthal sequences respectively displayed some activity above background levels (Figure 1D). Of these, 1,699 Denisovan (845 introgressed alleles, 854 AMH alleles) and 1,376 Neanderthal sequences (692 introgressed alleles, 684 AMH alleles) contained only a single allele, while the remainder contained multiple alleles in a haplotype construct (discussed further below). Finally, 22 of 249 measured random sequences (8.84%) were also deemed active, but these were less likely to be active across all replicates than test sequences or positive controls (Supplementary Figure 6). Activity ratios were moderately correlated across replicates, with activity of the positive control sequences showing the highest correlation of all sequence types (Supplementary Figure 7).

### Active introgressed alleles sequences display evidence of selective constraint

We then sought to characterise properties of active sequences to better understand what genomic features drive activity, as well as the features of functional archaic introgression. Approximately a quarter of active sequences were common to all five replicates (Supplementary Figure 6). We also observed higher sharing of Neanderthal active sequences between the two GM12878 replicates than sharing of Denisovan active sequences (145 Neanderthal sequences, 128 Denisovan sequences). More Denisovan active sequences were specific to the Papuan replicates (91) than Neanderthal active sequences (80). Furthermore, we did not see a bias towards either the modern or introgressed allele being active more often; approximately half of Denisovan and Neanderthal active sequences (49.7% and 50.3% respectively) contained the introgressed allele. Activity of the introgressed and modern alleles was clearly correlated (Figure 1D; Denisovan Pearson’s r = 0.667, p < 0.001; Neanderthal Pearson’s r = 0.669, p < 0.001).

We then asked whether allele frequency (AF) influences activity, as AF may be reflective of patterns of selection acting on functional archaic introgression. Because Neanderthal and Denisovan alleles segregating in genetically Papuan individuals have different population histories, we examined whether activity patterns change by decreasing AF decile for both introgressed and AMH alleles (Figure 1E, Supplementary Figure 8A), finding no significant association between AF decile and the percentage of active sequences within a decile (ANOVA p = 0.607). For Denisovan sequences, the first decile contains SNPs segregating at IAF ≥ 0.31, and at IAF ≥ 0.45 for Neanderthal sequences; the lack of an association between IAF and activity in general does suggest that introgression can rise to high frequencies even when functional.

However, we observed a significant association between distance to the nearest annotated transcription start site (TSS) and the likelihood of a sequence being active (Figure 1F, ANOVA p < 0.001). A much greater percentage of both modern and introgressed alleles were active when sequences were located within 1kb of the nearest TSS; however, this percentage is consistently lower for introgressed alleles than AMH ones. At increasing distances from the TSS the percentage of active sequences is approximately 10% regardless of introgression source or allelic state. Taken together, the relative high fraction of activity near TSS is likely to reflect general genome architecture (as well as the limitations of the MPRA system to capture small effects typically associated with distant enhancers), whereas the differences in activity between introgressed and modern alleles are suggestive of constraint against introgression in functional regions.

### Identifying differentially active introgression

Next, we tested for differential activity between the two alleles of active introgressed sequences. Throughout this section, we considered all 1,951 tested single-variant sequences with at least one active allele, and compared active alleles to their inactive counterparts when relevant. Using mpralm (Myint et al., 2019) we identified 180 sequences (95 Denisovan and 85 Neanderthal) with differential activity between the introgressed and non-introgressed alleles at an FDR threshold of 0.05 (9.22% of archaic SNPs tested for differential activity; Figures 2A, B, **Supplementary file 3**). Amongst these, the introgressed allele drives higher activity in 44 and 55 Denisovan and Neanderthal sequences respectively (46.3% and 64.7% respectively); and 27 sequences (12 Denisova, 15 Neanderthal) exceed log_2_ fold change values of 1. Of the differentially active Denisovan sequences, both alleles were active in 54 cases, while the remaining 41 showed activity in one allele only, while of the 85 differentially active Neanderthal sequences, 35 showed activity in both alleles, with 50 showing activity in one allele only.

**Figure 2.**
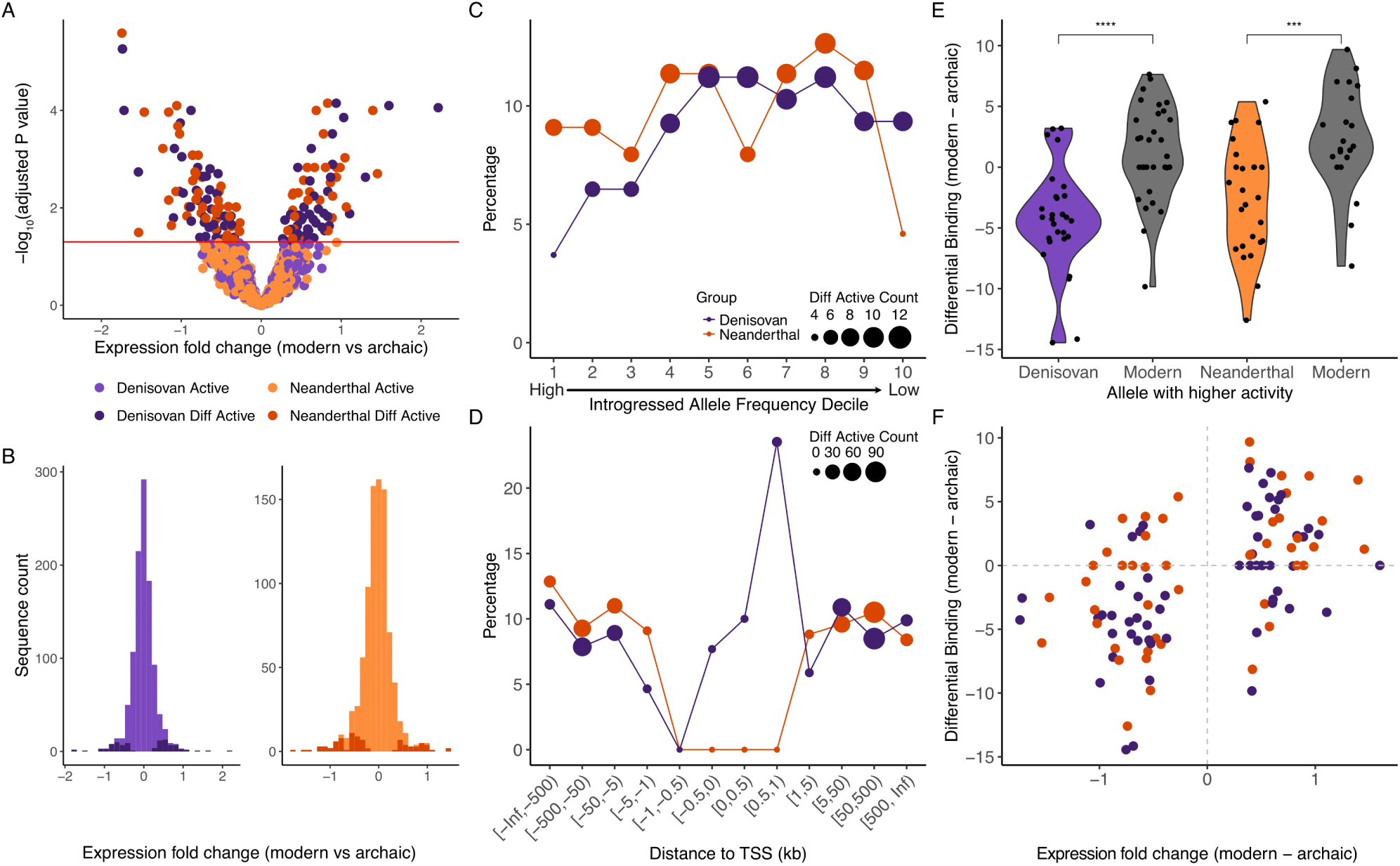
Identifying differentially active introgressed SNPs. A.) Volcano plot of results from mpralm (Myint et al., 2019). Red line represents an adjusted p-value = 0.05. B.) Distribution of expression fold change in activity driven by modern and archaic alleles for Denisovan and Neanderthal test sequences. C.) Percentage of differentially active SNPs across IAF deciles. D.) Percentage of differentially active SNPs in binned distances to the transcription start site (TSS). E.) Distribution of differential binding scores calculated with FIMO for differentially active SNPs, polarised by effect direction. Asterisks indicate Wilcoxon test p-values: *** = p *≤* 0.001,**** = p *≤* 0.0001 F.) Relationship between direction of differential activity and direction of differential binding score.

As above, we asked whether we could identify overall trends across differentially active alleles. Constraint on function at high IAF is clear when considering patterns of differential activity across deciles (Figure 2C). Denisovan and Neanderthal differentially active sequences display markedly different patterns, however. Differential activity increases from 3.7% in the highest IAF decile to an average of 10.4% for alleles in the bottom 60% of the data when considering Denisovan sequences, while IAF decile is less strongly associated with differential activity for introgressed Neanderthal sequences. However, the general paucity of introgressed alleles segregating at high allele frequencies (Supplementary Figure 3) means that the first IAF decile spans a large range of IAF values and obscures more fine-grained trends (Supplementary Figure 8B). No Neanderthal allele segregating at an IAF ≥ 0.55 is differentially active, while 1 Denisovan SNP with introgressed allele frequency between 0.55 and 0.65 is differentially active — rs143481175, is discussed further below and has the second strongest log_2_ FC in our data, -1.73, and an IAF of 0.64 in the (Jacobs et al., 2019) dataset. In general, we find no statistically significant correlation between log_2_ FC values and IAF for Denisovan (Spearman’s *ρ* = -0.19, p = 0.062) and Neanderthal (Spearman’s *ρ* = 0.056, p = 0.61) differentially active sequences. We also considered how distance to TSS of genes may influence differential activity (Figure 2D). No Neanderthal allele within 1kb of an annotated TSS is differentially active, which may reflect purifying selection. However, we did observe the highest percentage of differentially active Denisovan SNPs within 1kb downstream of the TSS, suggesting that introgression with a functional effect can be tolerated within regulatory elements.

Introgressed SNPs may alter enhancer activity through changes to transcription factor binding sites (TFBS; Vespasiani et al., 2022; Weyer and Pääbo, 2016; Findley et al., 2021). We used FIMO (Grant et al., 2011) to identify differential TF binding between the introgressed and non-introgressed alleles of a SNP. For differentially active SNPs, the direction of differential TF binding was consistent with the direction of differential activity; differentially active SNPs where the introgressed allele was more active had stronger TF binding to the introgressed allele than to the non-introgressed allele (Figure 2E; Denisovan Wilcoxon W = 730, p < 0.001; Neanderthal Wilcoxon W = 428, p < 0.001). In addition, differential binding score was correlated with estimated SNP effect size (mpralm logFC) for both Denisovan and Neanderthal differentially active SNPs (Figure 2F, Denisova: r = -0.499, p < 0.001; Neanderthal: r = -0.498, p < 0.001).

Taken together, these observations again suggest that generally, selective constraint acts to keep introgressed alleles with the ability to drive differential gene regulatory activity at low allele frequencies and away from genic TSS. Furthermore, the different patterns we observe across the two sources of introgression suggests different selection pressures acting on them.

### Assessing the interactive effects of introgressed alleles

Having quantified activity across single-variant sequences, we then turned to data derived from the 1,205 haplotype-like regions. The majority (87.5%) of these contained two variant sites, meaning four distinct sequences were needed to examine all possible allelic combinations (Figure 3A); smaller numbers of regions contained three, four and up to six variable sites (**Supplementary file 4**), requiring 2*^n^* sequences to capture all possible allelic combinations. Although all allele combinations were included in the originally synthesised oligonucleotide pool, only 6,537 haplotype-like alleles were considered for activity and differential activity testing.

**Figure 3.**
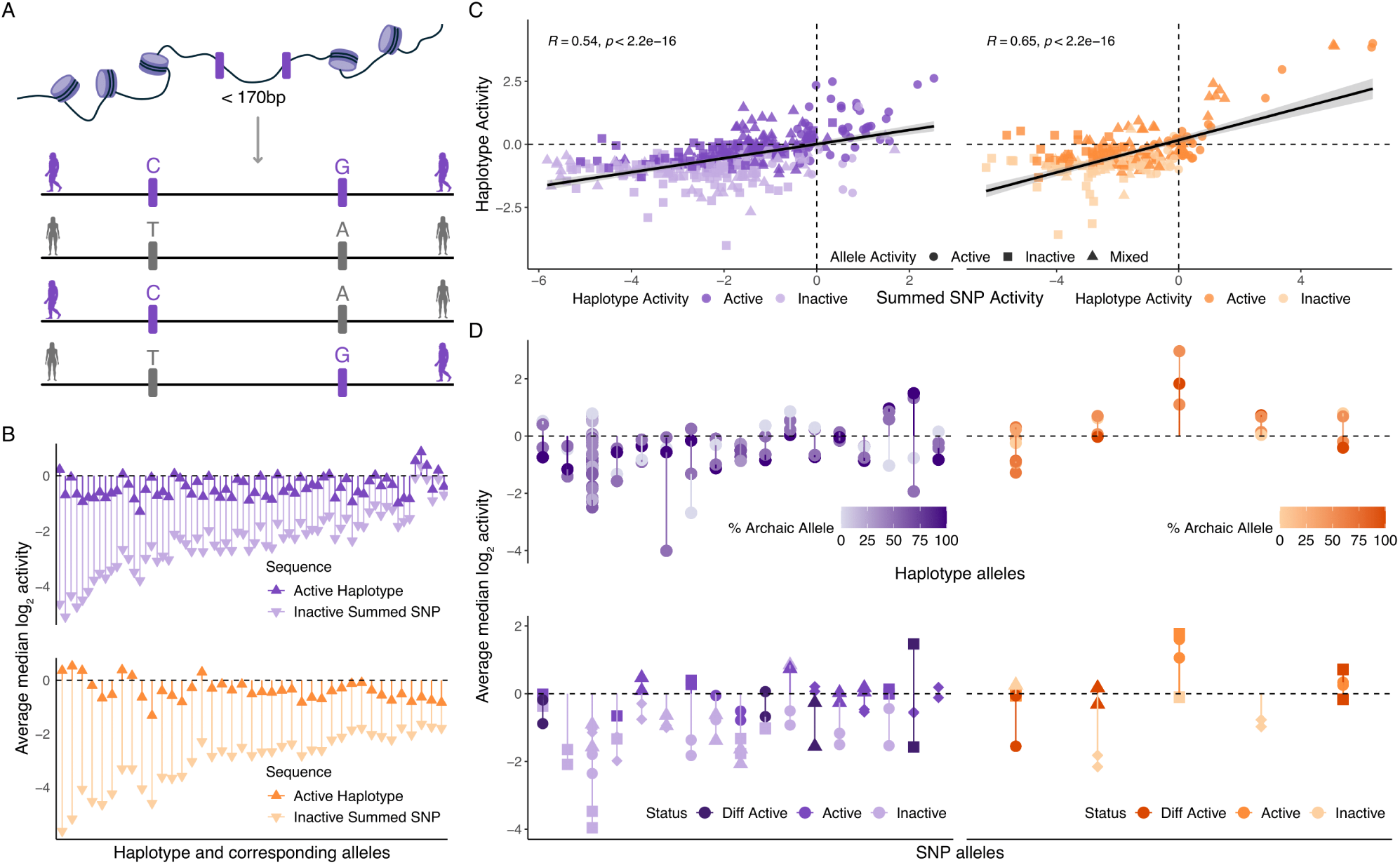
Testing of haplotype-like sequences. A.) Schematic of haplotype-like oligo design. B.) Activity of active haplotype-like sequences for which all corresponding single SNP sequences are inactive for Denisovan (top) and Neanderthal (bottom) sequences. Plots are ordered by decreasing difference between haplotype and summed SNP activity C.) Correlation between activity of haplotype-like sequences and the summed activity of the corresponding single SNP sequences for Denisovan (left) and Neanderthal (right) sequences. Mixed activity of summed single SNP sequences represents cases where the single SNP sequences for an given haplotype sequences were different in their activity status. D.) Activity of differentially active haplotypes (top row) and the activity of their corresponding single SNP sequences (bottom row) for Denisovan (left) and Neanderthal (right) sequences. Haplotype-like sequences are coloured by the percentage of introgressed alleles within the haplotype. Pairs of shapes for single variant sequences represents the two alleles of any given SNP.

Using the same approach as above, we classified 360 Denisovan haplotype-like sequences and 192 Neanderthal haplotype-like sequences as active, where “Denisovan” and “Neanderthal” indicate introgression source—not allelic state. These sequences represented 205 distinct genomic regions (126 Denisovan, 79 Neanderthal), and ranged from carrying exclusively introgressed alleles to exclusively non-introgressed alleles. There was no significant association between the fraction of introgressed alleles in a sequence and whether it was detected as active (Denisovan p = 0.224, Neanderthal p = 0.86).

Because all variants were also tested in single-variant sequences, we wanted to ask whether we could identify more complex interactions between variants when considering them jointly. 68 Denisovan haplotype-like sequences and 39 Neanderthal ones contained alleles that on their own did not show activity (Figure 3B) while the rest overlapped, at least partially, with some individual alleles identified as active above. Concordance between haplotype activity and the sum of their individual allelic activity was moderate (Denisovan Pearson’s r = 0.53, p <2.2 × 10^−16^; Neanderthal Pearson’s r = 0.64, p <2.2 × 10*^−^*^16^; Figure 3C). mpralm does not allow more than two conditions in differential activity testing. Thus, to test for differential activity of haplotype-like sequences, we performed an ANOVA on each haplotype locus, testing for differences in the activity of each allelic combination (**Supplementary file 5**). In this way we identified 22 haplotype sequences that showed differential activity at an FDR cutoff of 0.05 (Figure 3D). This consisted of 17 and 5 differentially active Denisovan and Neanderthal haplotypes, of which 13 and 2 respectively contained SNPs that on their own did not show differential activity.

### Predicting the effects of active and differentially active alleles

To identify potential phenotype effects of differentially active hits, we first used rGREAT (Gu and Hübschmann, 2023) to investigate annotations of the genes they are predicted to regulate. As expected given our variant selection strategy, genes near Denisovan active sequences were significantly associated with Gene Ontology (GO) biological processes related to immune cell differentiation and activation, while Neanderthal active sequences were associated with terms related to cell signalling and immune cell migration (**Supplementary file 6**). Denisovan ancestry active alleles were additionally associated with Human Phenotype Ontology terms related to facial, skin and skeletal phenotypes, while Neanderthal active sequences were associated with terms related to dental and skeletal morphology. While we did not observe any significant associations for genes predicted to be impacted by differentially active Denisovan sequences, those near differentially active Neanderthal sequences were associated with GO biological processes such as cell signalling, immune cell migration and lipid transport. When considering overlap with GTEx v8 (The GTEx Consortium, 2020) eQTLs, 16% of differentially active variants, 89.65% of which were Neanderthal, overlapped an eQTL in any GTEx tissue. This skew is unsurprising given the under-representation of non-European individuals in GTEx but serves as further validation of the power of our assay. Similarly, in 62% of cases we observed the same direction of effect across GTEx and our data. As in Tewhey et al., 2016 and Jagoda et al., 2022, concordance increased to 72.88% when the weakest one-third of eQTL results were removed.

We also examined individual sequences with strong evidence of differential activity between the introgressed and non-introgressed alleles, focusing on significant signals with large predicted fold changes. Genes near these hits include *IFIH1*, *TNFAIP3*, *UBS1* and *SNX9* amongst others. The most significant result was a Neanderthal variant, rs12464349 C>T, located in intron 11 of the *IFIH1* gene. The activity of the introgressed Neanderthal T allele was markedly higher than that of the non-introgressed allele (log_2_ FC = -1.743, p = 2.6 × 10^-6^), and highly reproducible across replicates (Figure 4A, **Supplementary file 3**). This site is located within an introgressed haplotype defined by Jacobs et al., 2019, which contains 46 testable variants of likely Neanderthal origin; of these 10 had at least one active allele in our data, but no others showed significant differential activity between the introgressed and non-introgressed allele (Figure 4B). *IFIH1* exhibits signals of adaptive introgression in Peruvians (Racimo et al., 2017; Witt et al., 2023) and individuals from the Bismarck Archipelago (Vernot et al., 2016). The frequency of the T allele is 0.47 in our data (Jacobs et al., 2019) and 0.035 in gnomAD v 4.1; allele frequency ranges from 0.47 to 0.93 in Native American populations from HGDP-CEPH, is between 0.27 and 0.10 in 10 different Chinese ethnic minority groups and does not rise above 0.10 anywhere else in the world (Supplementary Figure 9). While this pattern might suggest misattributed Denisovan ancestry, comparison against the three existing high-coverage Neanderthal genomes confirms this group as the likely source of the sequence (Supplementary Figure 10). A second haplotype, labelled “Haplotype 2” in Supplementary Figure 10, segregates in East Asian groups from the 1000 Genomes but not in the Papuan samples from Jacobs et al., 2019.

**Figure 4.**
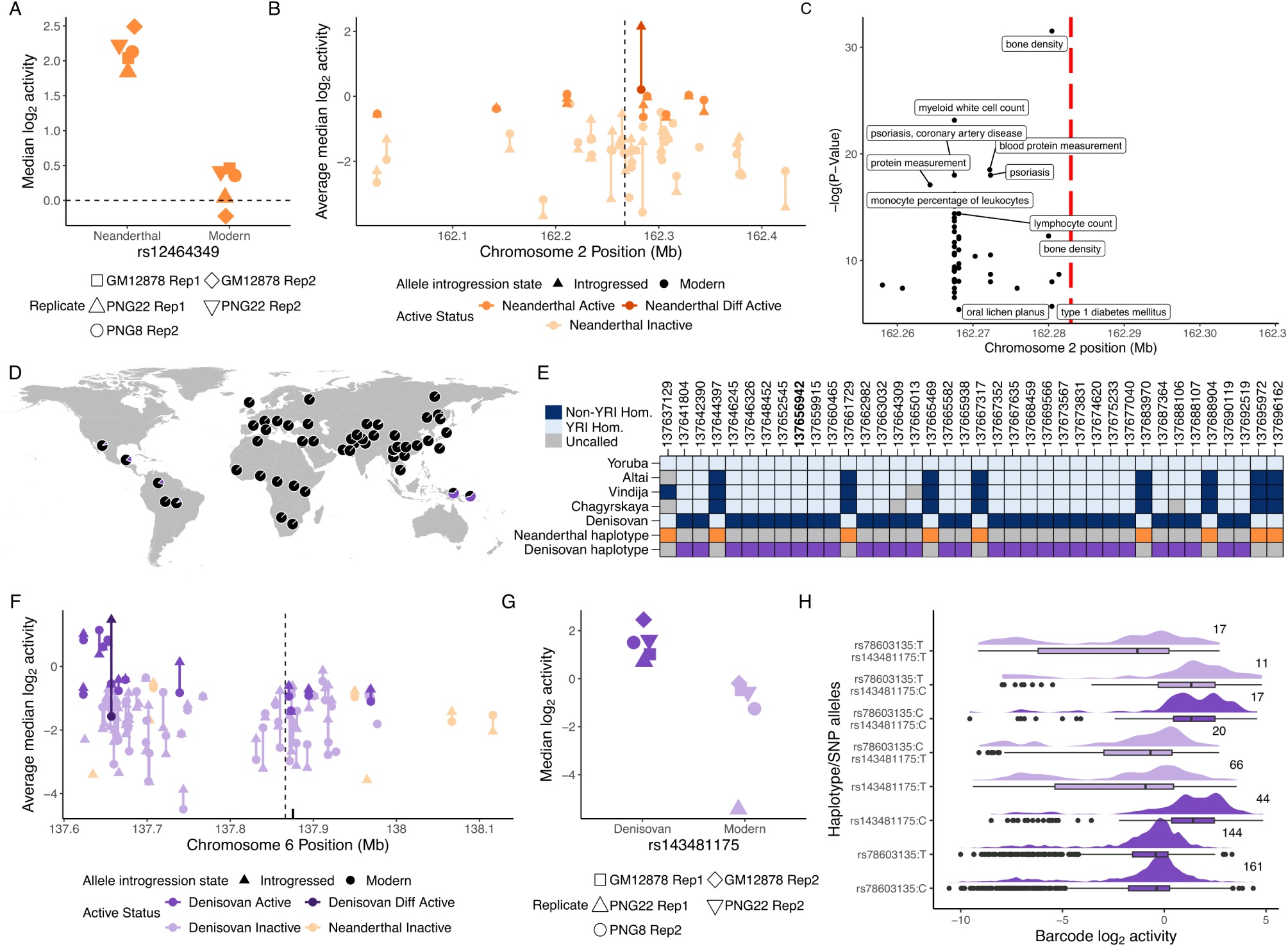
Differentially active SNPs are predicted to regulate immune genes. A.) Activity of rs12464349 alleles across replicates. B.) Allele activity difference for SNPs surrounding rs12464349. The transcription start site of *IFIH1* is indicated with a dashed line. C.) GWAS catalogue hits surrounding rs12464349 (indicated with a red line). D.) Introgressed allele frequency for rs143481175:C in the HGDP-CEPH populations, taken from gnomAD v 4.1. The pie centred over Port Moresby, Papua New Guinea, illustrates allele frequency in genetically Papuan individuals from Vespasiani et al., 2022. E.) Allelic states in archaic humans and the African Yoruba (YRI) population, together with the haplotype structure across archaic marker SNPs associated with archaic haplotypes surrounding rs143481175 (bolded position) in samples from Yermakovich et al., 2024. F.) Allele activity difference for SNPs surrounding rs143481175. The transcription start site of *TNFAIP3* is indicated with a dashed line. The positions of rs376205580 and rs141807543 are indicated with black lines. G.) Activity of rs143481175 alleles across replicates. H.) Distribution of barcode level activity for haplotype-like oligos containing rs143481175 and corresponding single-variant oligos across replicates. Numbers indicate the number of barcodes associated with each sequence.

*IFIH1* encodes MDA5, a cytoplasmic receptor that activates type I interferon signalling in response to dsRNA viral products. Gain of function mutations in *IFIH1* have been robustly linked to multiple interferonopathies (Rice et al., 2020), while loss of function variants have been repeatedly associated with increased risk of SLE (Graham et al., 2011) and inflammatory bowel disease (Adiliaghdam et al., 2022; Cananzi et al., 2021), but reduced risk of developing other autoimmune disorders, especially type 1 diabetes (Smyth et al., 2006; Nejentsev et al., 2009), suggesting a trade-off between sensitivity to antigens and autoimmunity (Gorman et al., 2017), akin to that described below for *TNFAIP3*. Although rs12464349 is not in the GWAS catalogue, likely due to its low global allele frequency, the surrounding region harbours multiple associations with immune phenotypes (Figure 4C).

We also identified two differentially active Denisovan SNPs predicted to regulate *IRF4* (rs932780686 C>G and rs764027350 C>G). Both SNPs are located on chromosome 6 downstream of the *IRF4* gene, and the Denisovan introgressed allele for both SNPs drive higher reporter gene expression than the modern alleles (rs932780686 log_2_ FC = -0.81, p = 4.37 × 10*^−^*^3^; rs764027350 logFC = -0.44, p = 0.0150; Supplementary Figure 11A-C). *IRF4* has been identified as a target of Denisovan adaptive introgression in Oceanian populations (Choin et al., 2021), and is critical for the development of lymphoid and myeloid cells, and regulates toll-like receptor signalling (Nam and Lim, 2016; Negishi et al., 2005), another known target of Denisovan introgression (Dannemann, Andrés, et al., 2016). In line with the role of *IRF4* in lymphocyte development, GWAS catalogue hits surrounding rs932780686 and rs764027350 were linked to immune cell phenotypes, including chronic lymphocytic leukemia and lymphocyte count, as well as autoimmune diseases, such as coeliac disease, rheumatoid arthritis and type 1 diabetes. Additionally, we see GWAS terms related to dermatological phenotypes, such as alopecia, freckles, hair colour and melanoma (Supplementary Figure 11D), consistent with the known association of Neanderthal introgression with skin, hair and eye pigmentation (reviewed by Reilly et al., 2022).

### A Denisovan variant predicted to impact *TNFAIP3* expression levels

Our most significant Denisovan differentially active sequence (log_2_ FC = -1.735, FDR-adjusted p = 5.46 × 10*^−^*^6^), and the second strongest signal in the data overall, was rs143481175 T>C, which has an IAF in Papuans = 0.64 (Jacobs et al., 2019) and a global AF in gnomAD v4.1 = 0.005991 (Figure 4D); this is the highest IAF amongst all differentially active sequences in our data. rs143481175 is located 162kb downstream of *OLIG3* and 210kb upstream of the *TNFAIP3* TSS. According to Roadmap Epigenomics data, rs143481175 falls within a predicted enhancer in the thymus, an immune tissue (ChromHMM state 7:Ehn; Kundaje et al., 2015). Furthermore, *TNFAIP3* has measurable expression in all GTEx v8 tissues, while *OLIG3* has no to very low expression (The GTEx Consortium, 2020). For these reasons we focus our analyses on the possible impact of rs143481175 on TNFAIP3 expression levels.

rs143481175 is located on an introgressed haplotype that spans roughly 120 kb on chromosome 6 (Figure 4E); the same Denisovan haplotype is present at a slightly lower mean IAF of 0.60, in the ≈ 300 samples from Yermakovich et al., 2024, which are a superset of the data in Jacobs et al., 2019 (Supplementary Figure 12). A second haplotype of likely Neanderthal ancestry also segregates in the same region at IAF of 0.12, while an additional Denisovan haplotype spans the *TNFAIP3* TSS and is also shown in Supplementary Figure 12. The high frequency of the Denisovan haplotype, together with the presence of a second Neanderthal haplotype in the same regions, provides a clear example of a genomic region significantly shaped by archaic introgression in Papuans. Across this entire region we tested 145 single-variant sequences for activity; 134 of these were Denisovan and the remaining 11 Neanderthal. 14 of these contained at least one active allele, all Denisovan, but only rs143481175 was found to be differentially active (Figure 4F). The Denisovan introgressed rs143481175:C allele reproducibly drove higher reporter gene expression than the modern allele (T), which was not classified as active (Figure 4G).

rs143481175 was also part of a haplotype-like sequences alongside rs78603135 C>T, which is located 56bp away from rs143481175 and has an IAF = 0.64 in our data. When tested on its own, both alleles of rs78603135 were active and we found no evidence of differential activity (log_2_ FC = 0.003, p = 0.988). Amongst the haplotype-like sequences, only the one containing rs78603135:C and rs143481175:C was labelled as active (Figure 4H). Despite having similar median activity to the single SNP oligos, the sequence containing containing rs78603135:T and rs143481175:C was not labelled as active, possibly due to having low barcode counts and a clearly bimodal signal distribution (Figure 4H).

*TNFAIP3* encodes the A20 protein, an ubiquitin modifying enzyme that negatively regulates TNF-NF-*κ*B signalling by regulating the ubiquitination of substrates in the NF-*κ*B signalling pathway (Wertz et al., 2004; Karri et al., 2024). Polymorphisms and mutations in *TNFAIP3* have been linked to autoimmune and inflammatory disease including SLE, rheumatoid arthritis and coeliac disease (reviewed in Ma and Malynn, 2012), and although rs143481175 is not in the GWAS catalogue, SNPs in the surrounding region have been associated with various autoimmune diseases including systemic lupus erythematosus (SLE), rheumatoid arthritis, coeliac disease and Sjogren syndrome (Supplementary Figure 13). The gene has been previously identified as harbouring strong signals of adaptive introgression in Papuans and Oceanians (Jacobs et al., 2019; Gittelman et al., 2016; Choin et al., 2021), and two missense mutations carried on a Denisovan introgressed haplotype (T108A, also known as rs376205580 A>G and I207L, also known as rs141807543 A>C) have been shown to have a slight, but significant effect on A20 activity (Zammit et al., 2019). Neither the Denisovan or modern human allele of these missense mutations were defined as active in our data, but this is to be expected, given that the experiment was designed to test for the potential to regulate gene expression.

## Discussion

Introgression from Neanderthals and Denisovans has shaped modern human genomes, but the exact molecular pathways and phenotypes impacted by it remain frequently unclear—especially when considering Denisovan introgression. Introgression is present in regulatory regions, particularly in those associated with immune cell types (Vespasiani et al., 2022; Dannemann, Andrés, et al., 2016; Gittelman et al., 2016; Nédélec et al., 2016; André et al., 2024; Yermakovich et al., 2024), suggesting that some introgressed variants may contribute to immune phenotypes through gene regulatory changes. At the same time, multiple lines of evidence point towards selection against Neanderthal (and likely Denisovan, but this has not been as thoroughly examined) introgression, likely driven by the high genetic load and low heterozygosity in the source populations (Harris and Nielsen, 2016; Sankararaman et al., 2016; Telis et al., 2020). Reconciling these two observations remains challenging.

Here, we have tested the regulatory potential of over 13,000 Denisovan and 11,000 Neanderthal variants segregating at moderate or high frequencies in individuals living in Papua New Guinea today, where Denisovan introgression levels are some of the highest globally. Only a small fraction (less than 10%) of tested alleles, regardless of ancestry, were capable of driving gene expression according to our assay. This number is lower than those reported in other MPRAs that have focused on human evolution (Jagoda et al., 2022; Uebbing et al., 2021; Weiss et al., 2021), which may reflect the biology of the types of variants being tested, but may also reflect differences in MPRA variant selection, experimental design, and analysis (Klein et al., 2020). Our estimate is also likely to be lower than the true value, as we only tested variants in a single cell type (see, for example, Tewhey et al., 2016; Weiss et al., 2021), and in the absence of any external stimulus. Nonetheless, we were able to identify introgressed and non-introgressed alleles capable of driving reporter gene expression, as well as SNPs where the two alleles drove significantly different reporter gene expression, which are strongly suggestive of functional potential *in vivo*. We also observed that the predicted direction of differential transcription factor binding was broadly consistent with the direction of observed differential activity in the MPRA, and the magnitude of both of these differences was well correlated, again supporting a functional role for these introgressed variants.

More broadly, we sought to understand the genomic forces that could impact the likelihood of an introgressed variant having functional consequences. Given conflicting reports of selection for and against introgression in regulatory regions, we examined whether distance to the nearest annotated transcription start sites was informative in this context. The percentage of active alleles did increase as distance to an annotated TSS decreased, but we also observed that a significantly higher percentage of non-introgressed alleles than introgressed alleles was consistently labelled as active near, but not far, from annotated TSS, suggesting some constraint on functional introgression at highly important functional regions. When considering differentially active alleles, we observed no differentially active Neanderthal SNPs within 1kb of annotated TSS, although we did observe instances of Denisovan differentially active SNPs in the same range, and similar patterns between the two introgression sources at greater distances. On the whole, these trends suggest that functional, and differentially functional, introgression can be tolerated within functional regulatory elements but is nonetheless subject to some degree of constraint.

Another example of forces that could help predict function is allele frequency: an introgressed variant can rise to high frequency both in the absence of positive selection—for example, through drift or hitch-hiking—or because of it. Because introgression is frequently detected at the level of haplotypes (even SNP-by-SNP methods group variants into haplotypes to control for incomplete lineage sorting), introgression catalogues are biased towards longer introgressed segments, which makes interpreting the results of selection scans on introgressed regions challenging. In our data, we found no significant association between allele frequency and the percentage of active sequences, suggesting that allele frequency alone is not enough to define or predict functional introgression. However, we did observe some evidence of constraint against differential activity at high IAF, particularly for Denisovan variants, and to some extent on Neanderthal variants, with no Neanderthal allele segregating at an IAF greater than 0.55 deemed differentially active. Yet despite this, the two strongest signals in our data came from alleles with high IAF: rs12464349, within an intron of *IFIH1* has an IAF of 0.47, and rs143481175, near *TNFAIP3*, has an IAF of 0.64 in the Jacobs et al., 2019 dataset, and one of the highest IAFs in the entire dataset. Notably, other variants in both of these genes have been implicated in evolutionary trade-offs between increased auto-immunity and decreased immune surveillance (Zammit et al., 2019; Gorman et al., 2017).

In our data, the Denisovan C allele at rs143481175 drove significantly higher levels of reporter gene expression than the modern human allele; suggesting it may increase expression levels of *TNFAIP3 in vivo* and possibly of A20 (the protein encoded by *TNFAIP3*). This increase may lead to increased downregulation of NF-*κ*B signalling pathways following immune activation, which could be beneficial in reducing chronic inflammation and autoimmunity—at the cost of decreased immune system responsiveness. Notably, the direction of this effect is opposite to that of two non-synonymous mutations introgressed from Denisovans (T108A and I207L) in *TNFAIP3*. Both of these mutations are common in Oceanic populations, including those that show otherwise low levels of introgression, but almost absent west of Wallace’s line (Zammit et al., 2019), suggesting positive selection drove their spread beyond genetically Papuan individuals. These variants give rise to a A20 protein with partial phosphorylation defects that lightly reduce the inhibitory activity of the protein, increasing immunity and resistance to coxsackievirus without causing spontaneous inflammatory disease that is associated with more severe loss of function. (Zammit et al., 2019). As such, the function of the A20 protein may be balanced by the effects of rs143481175 and the T108A/I207L alleles.

Unfortunately, the staggeringly poor representation of individuals carrying Denisovan introgression—and more broadly, of non-European genetic ancestries—in existing public genomic resources means further experimental validation of these results in orthogonal datasets is extremely challenging. For example, when looking for overlap between variants in the GWAS catalogue (Sollis et al., 2023) and our active SNPs, only 2 Denisovan and 5 Neanderthal SNPs were present in the GWAS catalogue. We were able to identify 274 active SNPs that are also eQTLs in GTEx v8 (The GTEx Consortium, 2020), with the majority of these being Neanderthal hits, as expected due to the disproportionate number of European donors in GTEx. These disparities have been well-documented before (e.g. Sirugo et al., 2019; Gallego Romero, Rodenberg, et al., 2024); both logistic and ethical challenges in generating these datasets outside the Global North (George et al., 2023) mean that they will be hard to redress in the short and middle term, at least at comparable scales to existing resources. Thus, MPRAs and other modalities of multiplex assays of variant effect (MAVEs), alongside computational predictors of variant effects, emerge as leading candidates to drive our understanding of the functional consequences not only of introgressed variants, but of genetic variation found in groups and individuals who are poorly represented in existing resources.

## Materials and Methods

### Ethics approvals and human samples

The two Papuan cell lines, PNG8 and PNG22, were established from healthy adult donors sampled at the University of Papua New Guinea, in Port Moresby, Papua New Guinea. Collection of blood was coordinated by Dr Christopher Kinipi (Director of Health Services at the University of Papua New Guinea) and approved by the Medical Research Advisory Committee of the National Department of Health of the Government of Papua New Guinea (permit number MRAC 16.21), by the University of Melbourne’s Human Research Ethics Committee (approvals 1851585.1 and 26981), and by the French Ethics Committees (Committees of Protection of Persons 25/21_3, n*^◦^*SI:21.01.21.42754). Permission to conduct research in Papua New Guinea was granted by the National Research Institute of Papua New Guinea (permit 99902292358), with full support from the School of Humanities and Social Sciences, University of Papua New Guinea. These approvals commit our team to following all ethical guidelines mandated by the government of Papua New Guinea. All individuals gave their full informed written consent to participate in the study.

### Variant selection and library design

To select variants for testing in the MPRA, we selected archaic variants segregating in genetically Papuan individuals from Jacobs et al., 2019 (for detailed information on defining archaic variants, see Vespasiani et al., 2022), if they were located in DNase I hypersensitivity sites or Tn5 transposase accessible regions in different immune cell types from the Roadmap Epigenomics Consortium (Kundaje et al., 2015), the ENCODE project (Thurman et al., 2012) and Calderon et al., 2019. We also included a set of archaic SNPs predicted to disrupt transcription factor binding site (Vespasiani et al., 2022). SNPs annotated with active chromatin states within monocytes, LCLs, B and T cells (Kundaje et al., 2015) were retained for testing. To prioritise variants of evolutionary and phenotypic importance, we additionally filtered for variants with a IAF of ≥ 0.15. Introgressed allele frequency was calculated as the number of observations of the introgressed allele divided by the number of observations for all alleles.

We additionally included two sets of control SNPs previously found to be active in MPRAs in LCLs (Tewhey et al., 2016) and in HepG2 cells (Arensbergen et al., 2019). For SNPs from Tewhey et al., 2016, we downloaded the full set of variants tested (https://www.cell.com/cms/10.1016/j.cell.2016.04.027/attachment/19be3b82-539c-4548-b522-cac927d98d55/mmc3.xlsx) and selected those with an rsid and absolute log2 allelic skew combined across the tested LCLs greater than 0.81. Where multiple alternative alleles were tested for the same rsid, we selected the allele with the maximum absolute log2 allelic skew. For SNPs from Arensbergen et al., 2019, we randomly selected 200 SNPs that were active in HepG2 cells, but not K562 cells.

Finally, we also designed a set of 300 random scrambled sequences to act as negative controls. For these random sequences, we confirmed the absence of any DNA motifs in HOCOMOCO v11 (Kulakovskiy et al., 2018) and JASPAR 2020 (Fornes et al., 2020) with FIMO (Grant et al., 2011) and the absence of matches to known human genomic DNA sequences with BLAT (Kent, 2002). We designed 170bp oligos centred on the selected SNPs, flanked by 15bp of an adapter sequence. Where two or more SNPs where located within 170bp of each other, we designed haplotype-like oligos containing all possible allelic combinations of the SNPs. We removed oligos containing SfiI and BsiWI restriction sites as these would be lost downstream in the cloning process. In total, our MPRA library included 61,812 oligos, testing a total of 25,869 introgressed SNPs (**Supplementary file 1**).

### Oligosynthesis and MPRA vector assembly

Oligos were synthesised by Twist Biosciences as 200bp single-stranded DNA sequences containing 170bp of genomic context centered on the variant of interest and 15bp of adapter sequence at either end (5’ACTGGCCGCTTGACG [170bp oligo] CACTGCGGCTCCTGC3’). We converted oligos to double-stranded DNA in a low cycle PCR (KAPA HiFi HotStart PCR Kit) with primers designed to bind to the adapter sequences (F_adapter and R_adapter, **Supplementary file 7**). We next added 15bp random barcodes, SfiI and BsiWI restriction sites, HiFi cloning sites and a buffer sequence to the oligos in another PCR reaction (NEB Q5 High-Fidelity 2x Master Mix; F_PCR and Barcoding_R, **Supplementary file 7**). Barcoded oligos and pMPRA1 plasmid were digested with SfiI (NEB) and the digested oligos ligated into the pMPRA1 plasmid with T4 ligase (NEB). pMPRA1 was a gift from Tarjei Mikkelsen (Addgene plasmid # 49349, Melnikov, Zhang, et al., 2014). The resulting plasmid library was transformed into 10-beta electrocompetent E. coli (NEB) using a BioRad GenePulsar (2.0 kV, 200 Ω and 25 *µ*F) and allowed to grow overnight in five flasks containing 500ml LB-SeaPrep agarose (Lonza) culture (Elsaesser and Paysan, 2004) and ampicillin at a final concentration of 0.1mg/ml. Colonies were pelleted via centrifugation and giga-prepped (Zymo). To associate each barcode with its corresponding oligo, we amplified a fragment containing the oligo and barcode from each plasmid with PCR (NEBNext Ultra II Q5 Master Mix). Fragments were sequenced by Azenta with 150bp paired end reads on Illumina Novaseq. We used the MPRAmatch pipeline (https://github.com/tewhey-lab/MPRA_oligo_barcode_pipeline, adapted with custom scripts)) to associate barcodes with oligos (**Supplementary file 8**).

The plasmid library was digested with BsiWI (NEB) and GFP (IDT) was cloned into the plasmid vector with HiFi assembly (NEBuilder HiFi DNA Assembly MasterMix). The final plasmid library was transformed into 10-beta electrocompetent E. coli (NEB) using a BioRad GenePulsar (2.0 kV, 200 Ω and 25 *µ*F) and allowed to grow overnight in four flasks containing 500ml LB with ampicillin (final concentration = 0.1mg/ml). Colonies were pelleted via centrifugation and giga-prepped (Zymo). The concentration of the resulting plasmid preparation was increased with ethanol precipitation.

### Cell culture and transfection

Three lymphoblastoid cells lines, GM12878 (Coriell), PNG22 and PNG8 were grown in suspension in RPMI (Gibco) supplemented with 10% FBS (Scientifix), 1% NEAA (Gibco) and 1% Glutamax (Gibco), maintaining a density of 0.2-0.5 × 10^6^ cells per mL until there was a total of 120 × 10^6^ cells per line, except for GM12878 replicate 1, where we grew 160 × 10^6^ cells. The day prior to transfection with a Nucleofector II (Lonza), cells were seeded at a concentration of 0.5×10^6^. The day of transfection, cells were counted using a Countess 3 (Thermo Fisher) and pelleted at 500xg for 5 minutes. Cells were resuspended in supplemented Nucleofection solution V. The plasmid library was added to this solution, and 100*µ*l cell-DNA suspension was transferred to a nucleofection cuvette (each cuvette contained 5 x 10^6^ cells and 40*µ*g plasmid library). The cuvette was inserted into the Nucleofector II device and program X-001 applied. Immediately after, 500*µ*l of prewarmed culture medium was added to the cuvette. The nucleofected cell suspension was then transferred to a T75 flask and incubated for 24 hours. This process was repeated until all cell-DNA suspension has been transfected (12-16 reactions per replicate). 24 hours post-transfection, cells were pelleted for 5 minutes at 500xg. The media was removed and cell pellets resuspended in 2ml DPBS. 90% of the suspension was taken for RNA extraction and 2 x 5% was taken for DNA extraction. Cells were spun down again at 500xg for 5 minutes. The DPBS was removed and the cell pellets frozen at -80°C for storage. Two replicates were performed per cell line, with 60 × 10^6^ cells for all replicates except GM12878 replicate 1, where we transfected 80 × 10^6^ cells.

### DNA/RNA extraction and cDNA synthesis

Plasmid DNA (pDNA) was extracted from the cell pellets using Qiagen’s DNeasy Blood and Tissue Kit following manufacturer’s protocols, performing an optional digestion with RNase A (Qiagen). RNA was extracted from the cell pellets using Qiagen’s RNeasy Midi kit, performing an optional on-column DNase digestion (Qiagen). RNA extractions were treated with Turbo DNase (Thermo Fisher) to remove any DNA carryover. First-strand cDNA was synthesised with Superscript IV (Thermo Fisher) using a primer designed to bind to the buffer sequence (reverse_transcription_primer, **Supplementary file 7**). To quantify the number of cycles required for library amplification, we performed qPCR (NEBNext Ultra II Q5 master mix 2x) for both pDNA and first-strand cDNA using primers binding to GFP and the buffer sequence (GFP_F and Buffer_sequence_R, **Supplementary file 7**). Using the Ct values determined from qPCR, the barcode region of the cDNA and pDNA libraries was amplified with PCR (NEBNext Ultra II Q5 master mix 2x; GFP_F and Buffer_sequence_R, **Supplementary file 7**). PCR products were cleaned using Qiagen’s MinElute kit, and checked for product size and concentration with Agilent Tapestation. Samples were intially sequenced to 20M paired end reads on Illumina NovaSeq by Azenta. After confirming the quality of the sequencing library, the samples were resequenced to 240M paired end reads. An additional replicate (PNG8 rep1) did not have enough cDNA to be sequenced, so it was ommited from all analyses.

### Data analysis

Adapter sequences were removed from reads from both rounds of sequencing with Cutadapt v4.8 (Martin, 2011). Paired end 150bp reads were merged with Flash v2.2 (flags: -r 150 -s 30 -t 25) (Magoč and Salzberg, 2011) and reads from the two rounds of sequencing merged. Reads were passed to the MPRAcount pipeline (https://github.com/tewhey-lab/MPRA_oligo_barcode_ pipeline, adapted with custom scripts) to count the number of each barcode in the cDNA and pDNA libraries (Supplementary Figure 14). Following methods from Uebbing et al., 2021, cDNA and pDNA counts were normalised by library size and log2 transformed (**Supplementary file 9**). Barcodes with an average pDNA log2CPM < −5 across replicates were removed (Supplementary Figure 15). We also removed oligos for which there were fewer than 10 associated barcodes. For each barcode, we calculate its ’activity’ as the ratio of cDNA counts over pDNA counts. To define active sequences, we compare the activity distribution of all barcodes for an oligo against the mean activity per replicate using a one-sided *t* test. P-values were corrected with Benjamini-Hochberg multiple testing correction, and we define a fragment as active if it has a corrected p-value < 0.05 in at least two replicates. Sequences were defined as European specific if their corrected p-value was < 0.05 in both European replicates and > 0.05 in all three Papuan replicates. Sequences were defined as Papuan specific if their corrected p-value was < 0.05 in at least two of the three of the Papuan replicates and > 0.05 in both European replicates.

All sequences with at least one active allele were then tested for differential activity. Where relevant, we compare active alleles to their inactive counterparts. We take two approaches to differential activity testing. First, to test for differential activity between two alleles of a SNP, we used the mpra v1.26 R package (Myint et al., 2019). We used the *mpralm()* command, which is based on the voom framework (Law et al., 2014), to test for differential activity between the introgressed and modern allele for Denisovan and Neanderthal sequences. cDNA and pDNA counts for each oligo were summed across barcodes and the aggregated counts passed to the command

mpralm_sum <− mpralm ( o b j ect = mpraSet_sum , design = design_sum, block = block_vector_sum , aggregate = “ none “ , normalize = TRUE, model_type = “ corr_groups “ , p l o t = TRUE)

We define sequences as differentially active between the introgressed and modern alleles if it has a Benjamini-Hochberg adjusted p-value <0.05. To test for differential activity of haplotype-like sequences, we apply ANOVA to test for differences in activity of each oligo per replicate. Haplotype-like oligos were defined as differentially active if they had a Benjamini-Hochberg adjusted p-value <0.05 in at least two replicates. Activity testing and subsequent analysis was performed in R v.4.4.0.

### SNP annotation

To assess whether allele frequency influences activity, allele frequencies were calculated as in Vespasiani et al., 2022. For fair comparison of the frequency of introgressed and modern alleles, for each introgression source (Denisovan/Neanderthal) and allelic state (introgressed/modern), we divide sequences into allele frequency deciles, and calculate the percentage of active alleles within each decile. Similarly, to assess whether allele frequency influences differential activity, we divide sequences into introgressed allele frequency deciles, and calculate the percentage of differentially active sequences within in decile.

To assess how distance to TSS influences activity, we used the *getRegionGeneAssociations* command in rGREAT v2.6 (Gu and Hübschmann, 2023) to link sequences to genes they putatively regulate. We run rGREAT on our active and differentially active hits using hg38 coordinates (converted from hg19 coordinates with rtracklayer v1.64 (Lawrence et al., 2009)), separately for Denisovan and Neanderthal hits, with the *great()* command, specifying mode = “twoClosest” and tss_source = “Gencode_v40”. We binned distances to the TSS, and calculated the percentage of active and differentially active sequences within each distance bin.

To identify enriched Gene Ontology and Human Phenotype Ontology terms, we used rGREAT to test for enrichment of active and differentially active hits in MSigDB GO and HPO gene sets, using the set of sequences tested for activity and differential activity as background sets. Enrichments were considered significant at FDR <0.05.

To assess the disruption of TFBS, all sequences in the MPRA library were first trimmed to the middle 40bp. The trimmed list of sequences was passed to FIMO (Grant et al., 2011) to query the trimmed sequences for motifs in the HOCOMOCO v11 core collection (Kulakovskiy et al., 2018), specifying to only include sequences with p-value <0.0001 in output. We filtered for Denisovan and Neanderthal sequences where one allele has a p-value <10^-4^ and the other has a p-value <10^-3^. For these sequences, we calculate a differential binding score between the introgressed and the modern allele by taking the difference between FIMO binding scores for the two alleles. Where a SNP was predicted to disrupt multiple TFBS, we take the motif with the largest absolute differential binding score.

To search for SNPs in the GWAS catalogue, we used v1.0.2 of the GWAS catalogue (downloaded from https://www.ebi.ac.uk/gwas/docs/file-downloads). We searched for matching rsIDs between our MPRA SNPs and the GWAS catalogue. To define GWAS SNPs in the regions surrounding our differentially active hits, we searched the GWAS catalogue for SNPs within 50kb of each differentially active SNP.

To search for eQTLs overlapping and surrounding our hits, we downloaded GTEx v8 (The GTEx Consortium, 2020) significant variant-gene pairs for all tissues (https://gtexportal.org/home/downloads/adult-gtex/qtl). We merged eQTLs for all tissues, and overlapped these with the set of variants tested in our MPRA by chromosome, hg38 coordinates, the reference and alternative allele. To examine concordance between our data and GTEx, we transformed the mpralm effect size to reflect the effect of the alternative allele against the reference allele, and compare the direction of this effect to the slope of the eQTL. To assess concordance after removing weak effects, we removed the smallest one-third of absolute effect sizes, and recalculated concordance.

### Data access

Raw sequencing reads have been deposited to ENA under accession number PRJEB88773. Code for all analyses is available here: https://gitlab.unimelb.edu.au/igr-lab/PNG_introgression

## Acknowledgements

This work was supported by an award from the Leakey Foundation and by Australian Research Council Discovery Project DP200101552, both to IGR, by the Marsden Fund of the Royal Society of New Zealand (20-MAU-017) to MPC and IGR, by French National Research Agency (ANR) (grant PAPUAEVOL ANR-20CE12-0003-01 to FXR, and by Estonian Research Council grant TK (TK214) to MD. St Vincent’s Institute acknowledges the infrastructure support it receives from the National Health and Medical Research Council Independent Research Institutes Infrastructure Support Program and from the Victorian Government through its Operational Infrastructure Support Program. The funders had no role in study design, data collection and analysis, decision to publish, or preparation of the manuscript. We thank other members of the Gallego Romero lab, Pradiptajati Kusuma and Mait Metspalu for valuable comments on the manuscript, and sample donors at the University of Papua New Guinea for their contribution to this work.

## Author contributions

- Conceptualization: IGR, DMV
- Formal analysis: MC, DMV, MD
- Funding Acquisition: IGR, FXR
- Data Curation: MC, MF, DY
- Investigation: MC, DMV
- Methodology: MC, DMV, LEC, NS
- Resources: ML, CK, FXR, NB DY, MD
- Supervision: IGR
- Visualization: MC, MD
- Writing – Original Draft Preparation: MC, IGR
- Writing – Review & Editing: All

## Supplementary Figures

**Supplementary Figure 1.**
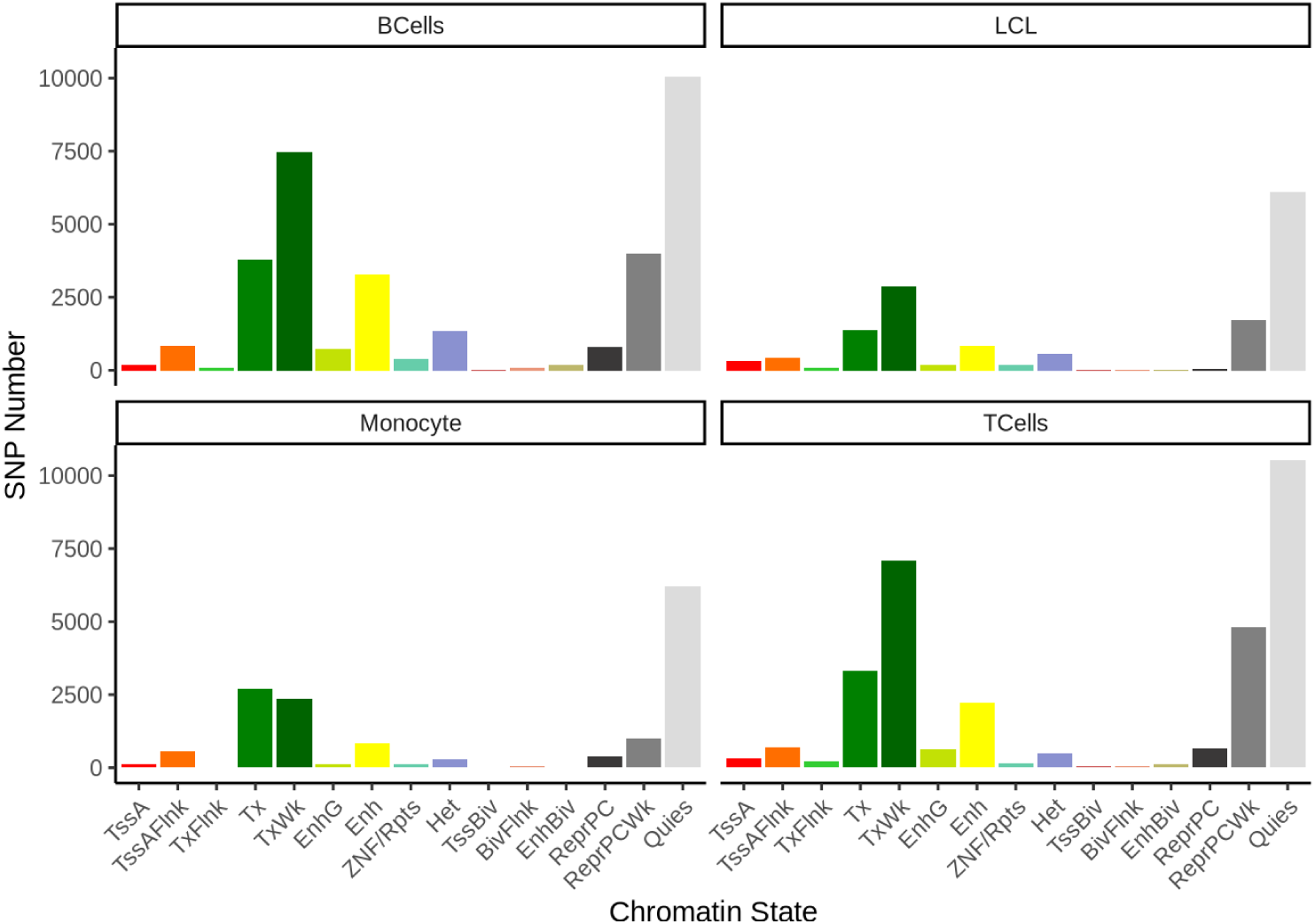
Number of Denisovan SNPs in initial library in chromHMM chromatin states across immune cell types.

**Supplementary Figure 2.**
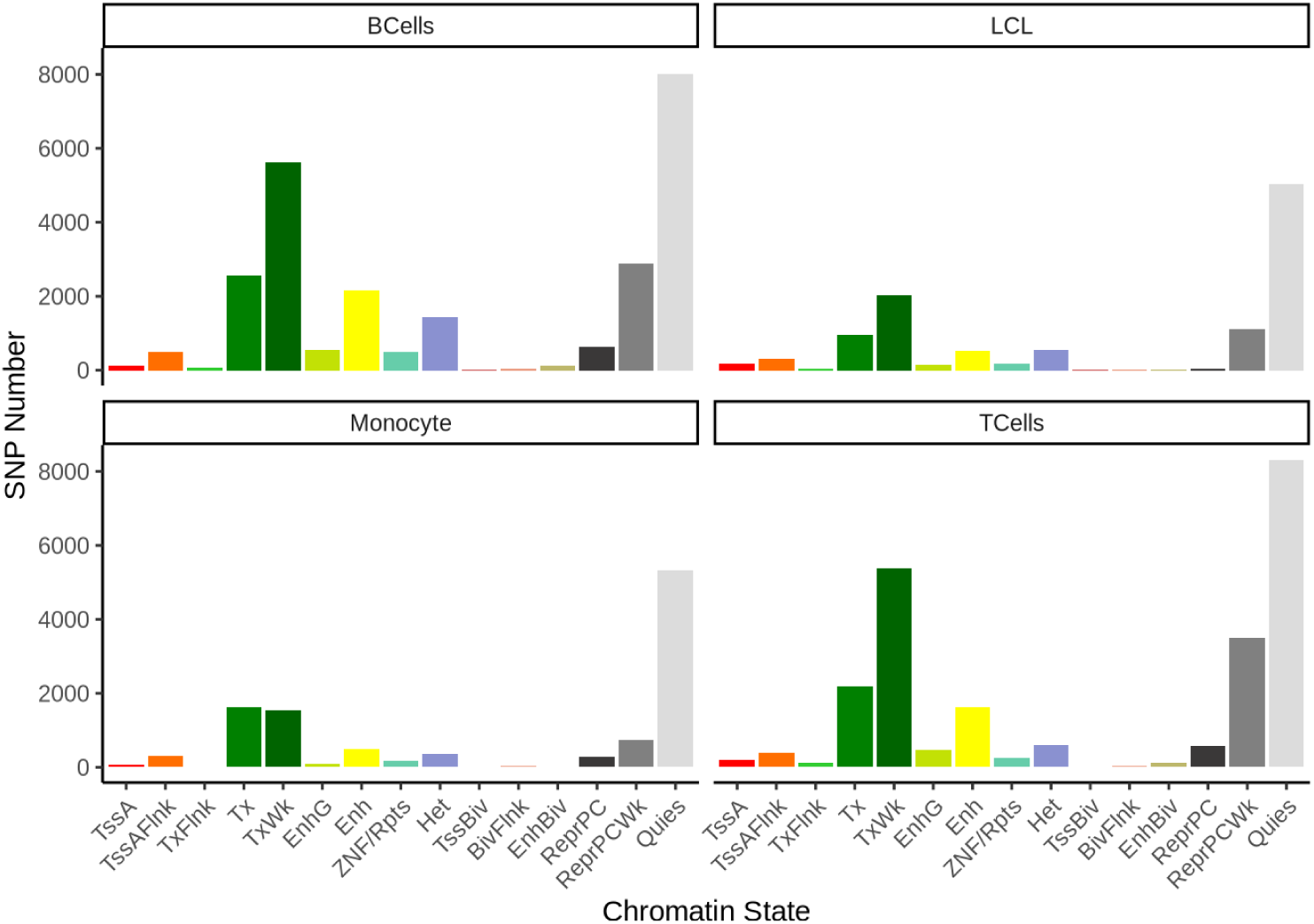
Number of Neanderthal SNPs in initial library in chromHMM chromatin states across immune cell types.

**Supplementary Figure 3.**
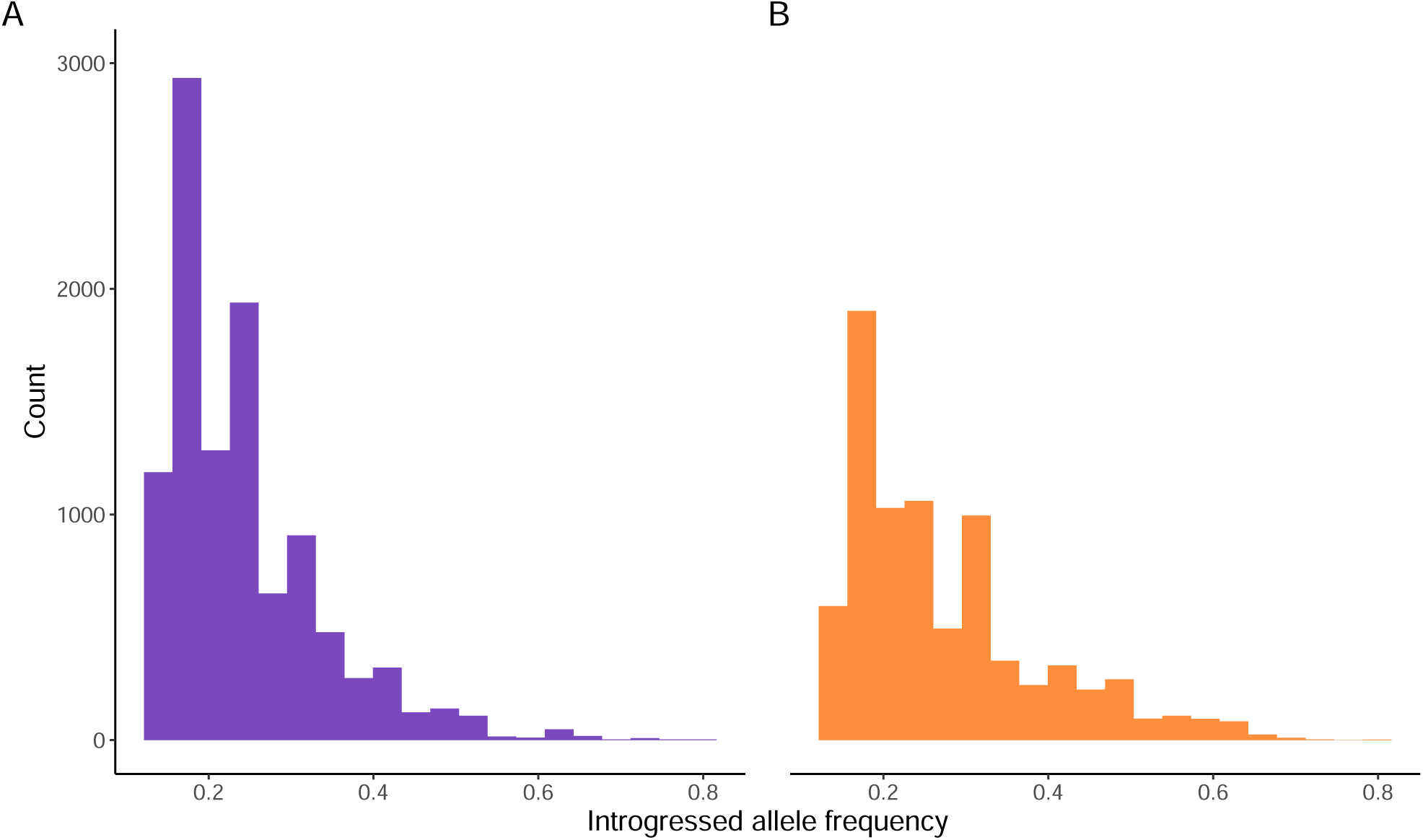
Distribution of IAF for A) Denisovan and B) Neanderthal measured alleles.

**Supplementary Figure 4.**
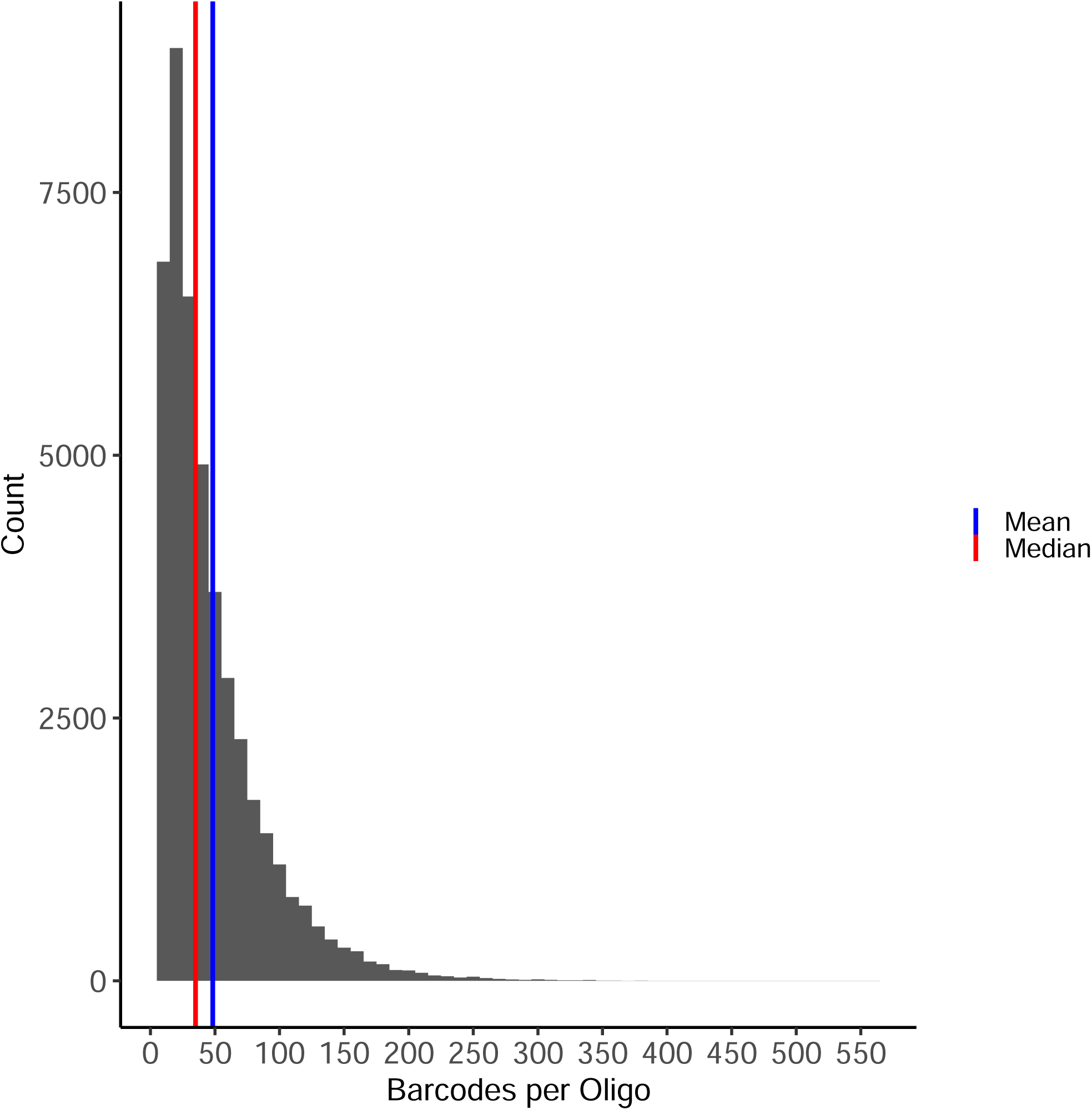
Distribution of barcodes per oligo post filtering.

**Supplementary Figure 5.**
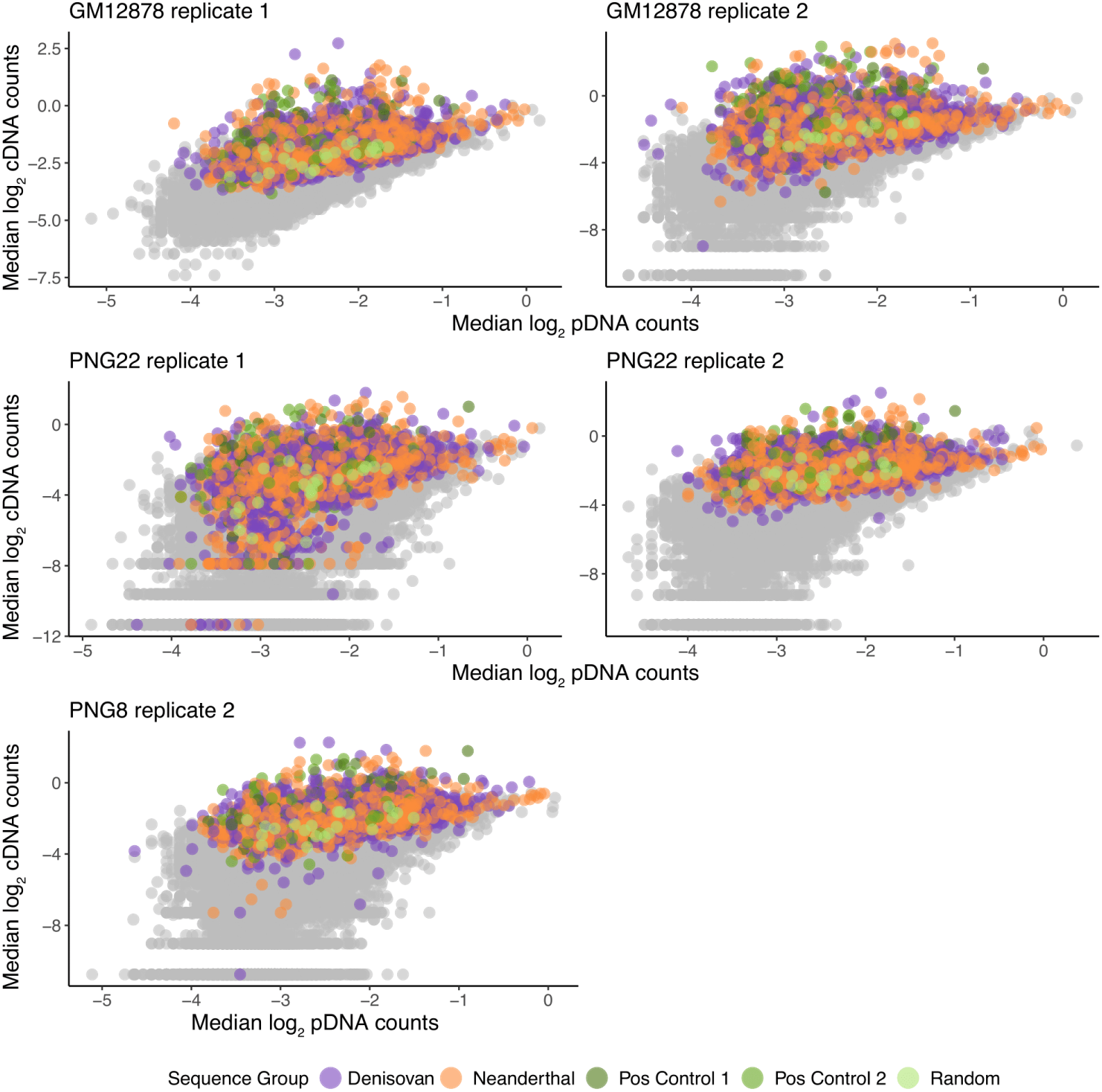
Scatterplots of cDNA and pDNA counts per replicate. Active sequences are highlighted in colour. Positive control 1 and 2 sequences were previously found to be active by Tewhey et al., 2016 and Arensbergen et al., 2019, respectively.

**Supplementary Figure 6.**
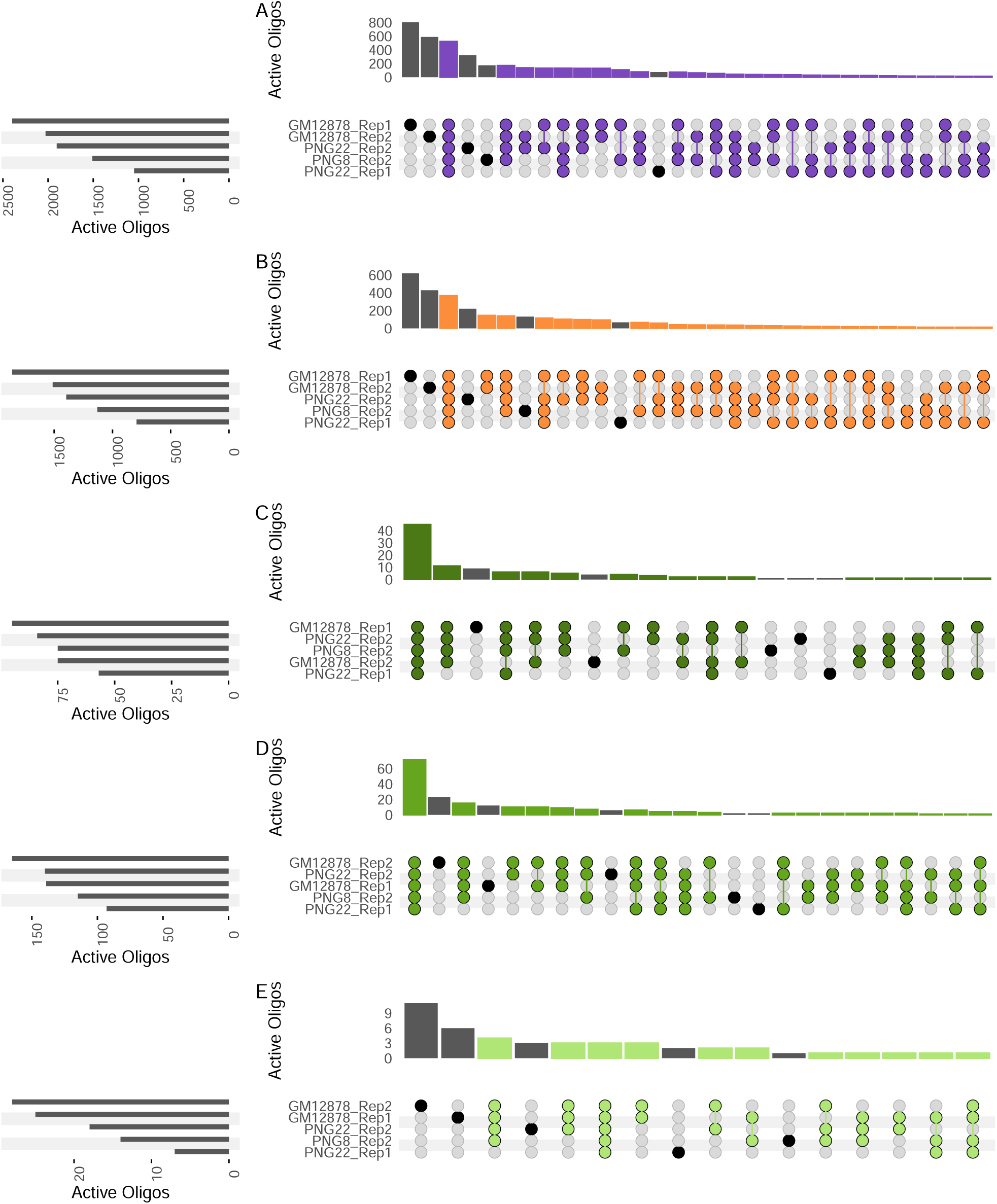
Sharing of activity across replicates for A) Denisovan, B) Neanderthal, C) positive controls from Tewhey et al., 2016, D) positive controls from Arensbergen et al., 2019 and E) random scrambled sequences. Sequences were labelled as active if active in at least two replicates.

**Supplementary Figure 7.**
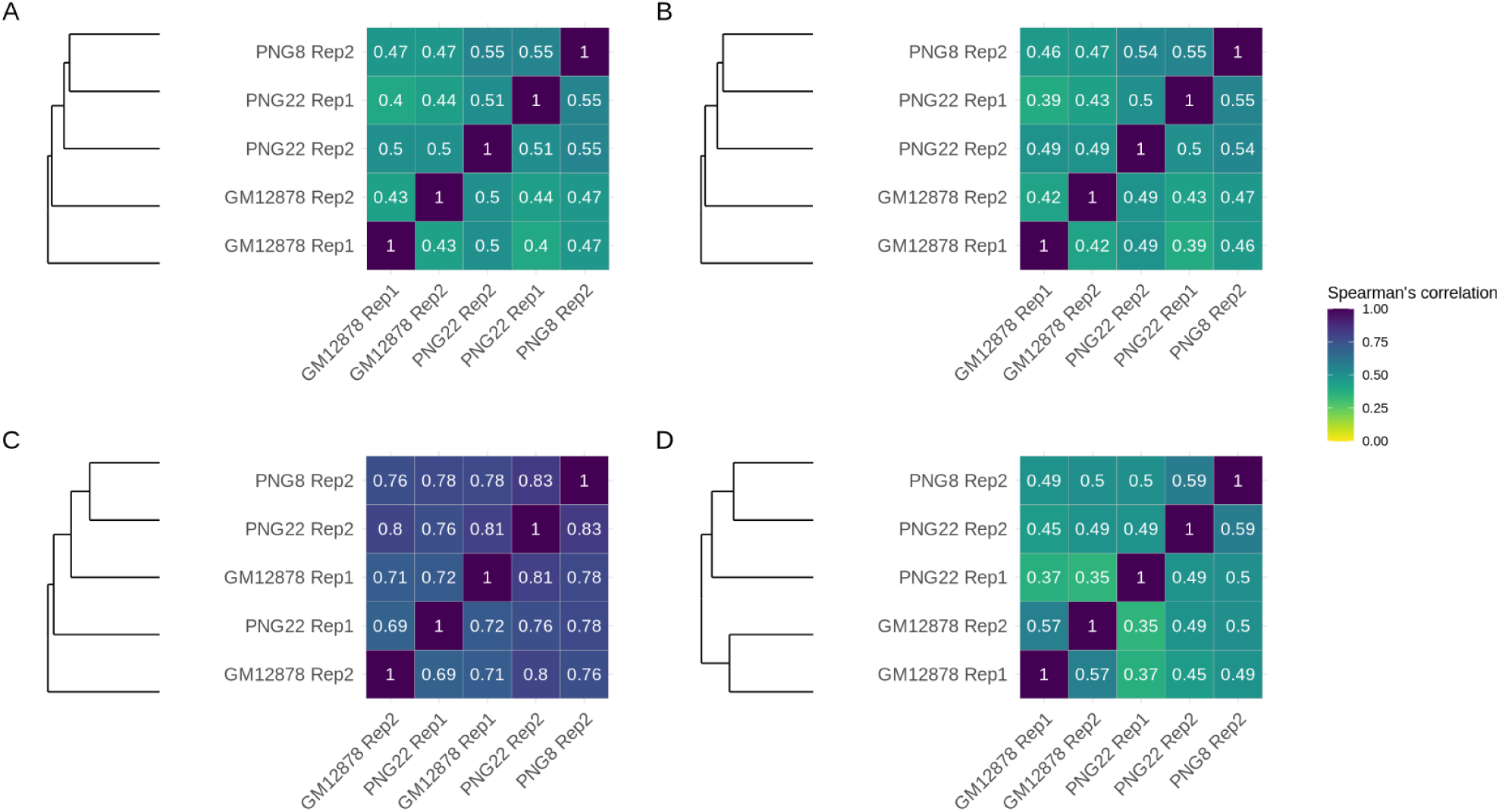
Spearman’s correlation of activity across replicates for A) all sequences, B) archaic sequences, C) positive control sequences and D) random scrambled sequences.

**Supplementary Figure 8.**
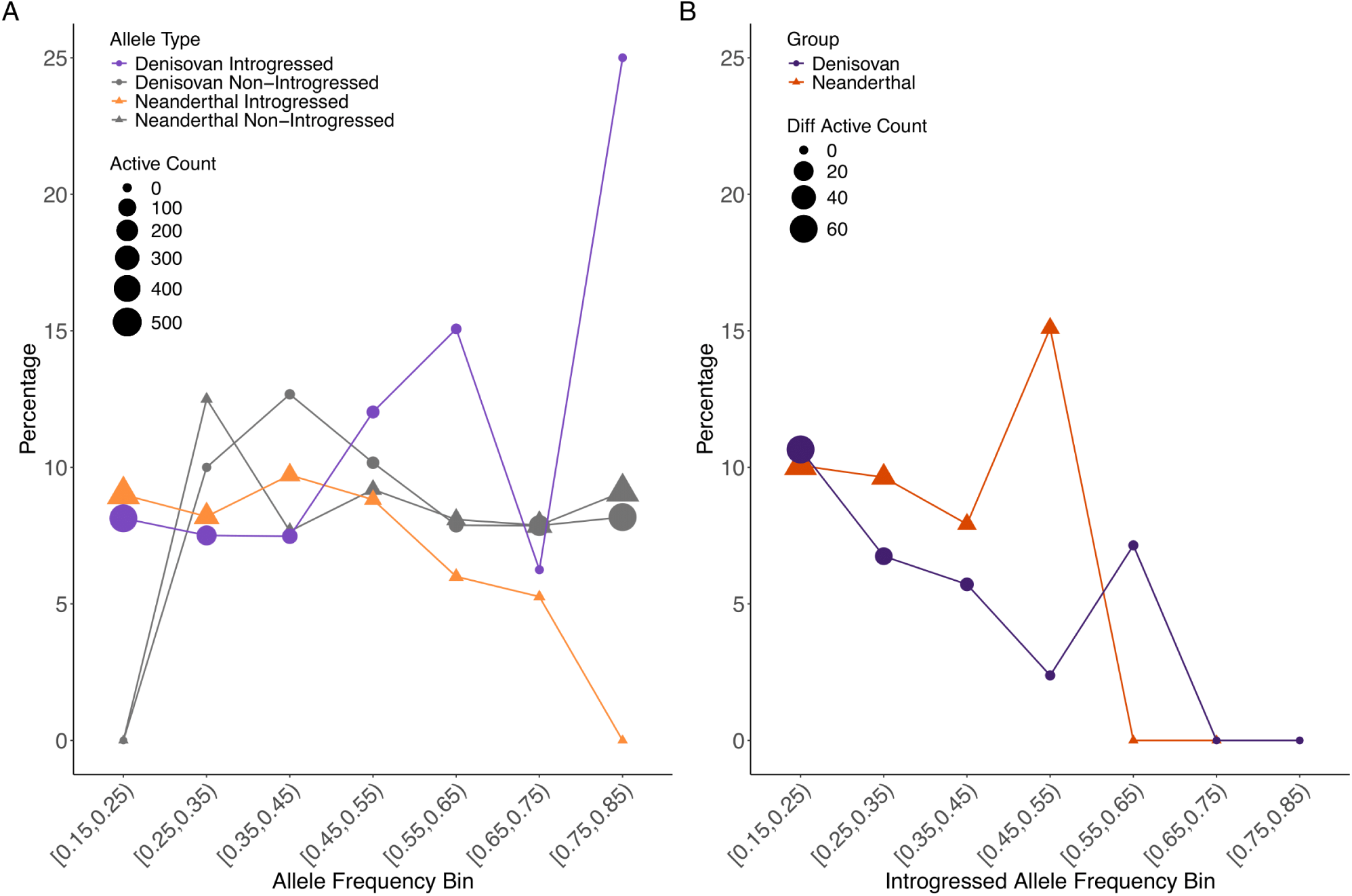
Percentage A) active by allele frequency bins and B) differentially active hits by introgressed allele frequency bins.

**Supplementary Figure 9.**
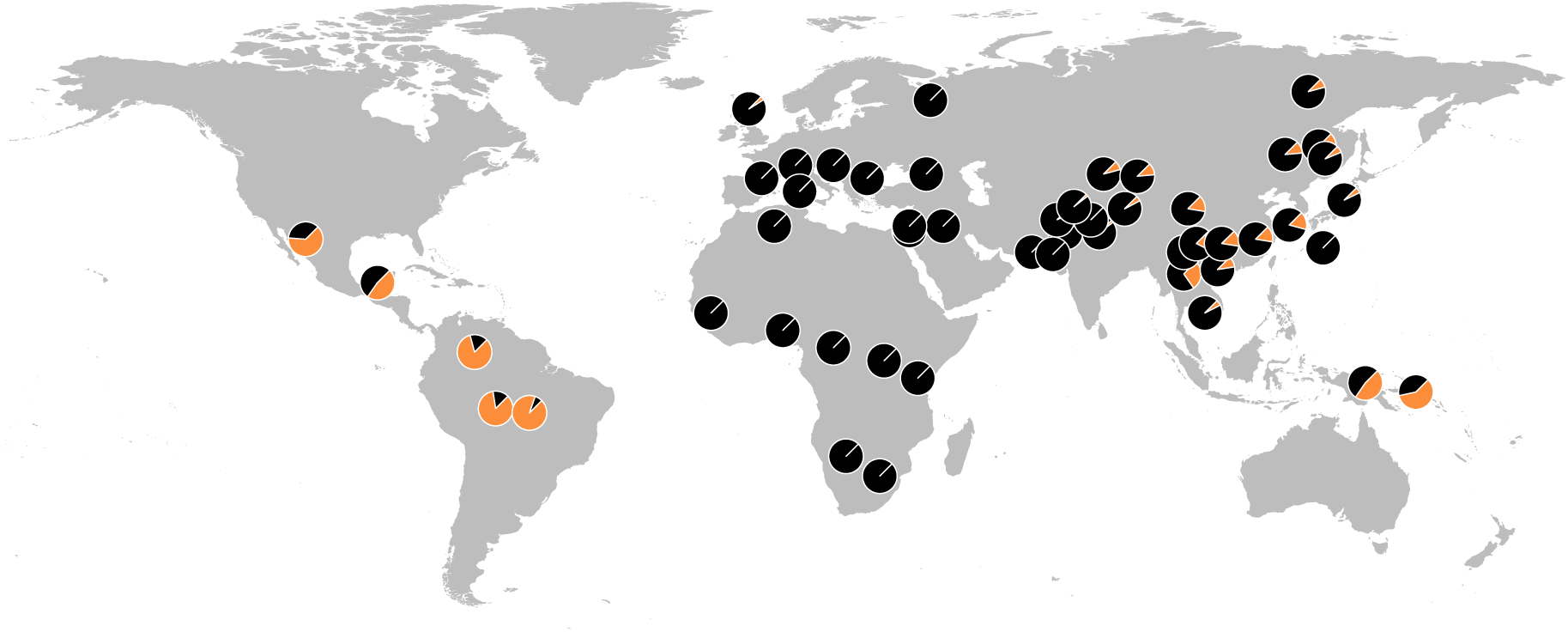
IAF for rs12464349:T in the HGDP-CEPH populations, taken from gnomAD v 4.1. The pie centred over Port Moresby, Papua New Guinea, reports allele frequency in genetically Papuan individuals included in Vespasiani et al., 2022 instead.

**Supplementary Figure 10.**
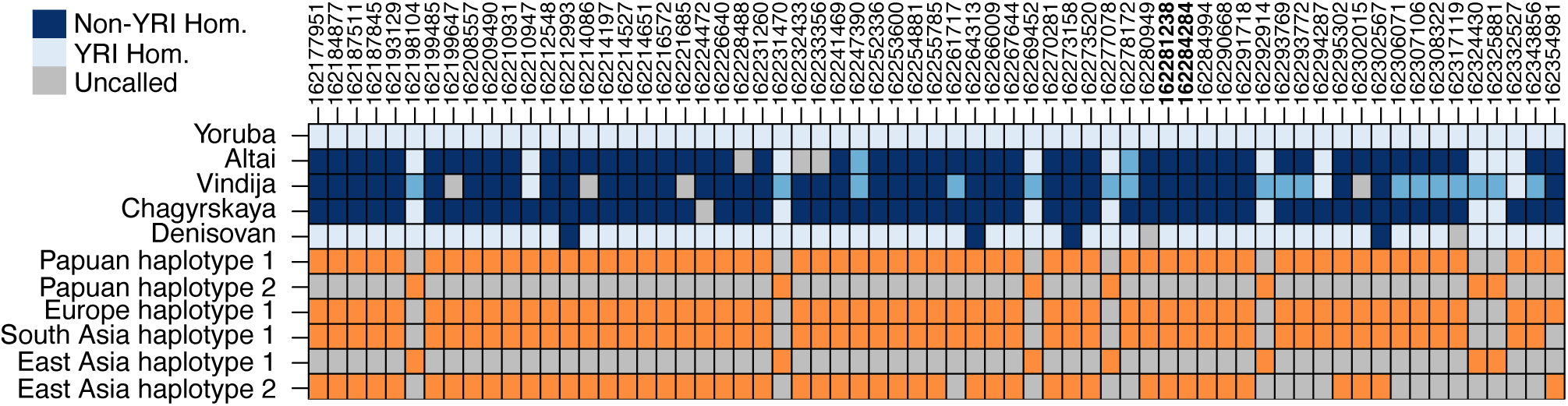
Two differentially active SNPs predicted to regulate *IFR4*. A). Activity of rs932780686 alleles across replicates. B). Activity of rs764027350 alleles across replicates. C.) Allele activity difference for SNPs surrounding rs932780686 and rs764027350. The transcription start site of *IRF4* is indicated with a dashed line. D.) GWAS catalogue hits surrounding rs932780686 (indicated with a dashed line) and rs764027350 (indicated with a solid line).

**Supplementary Figure 11.**
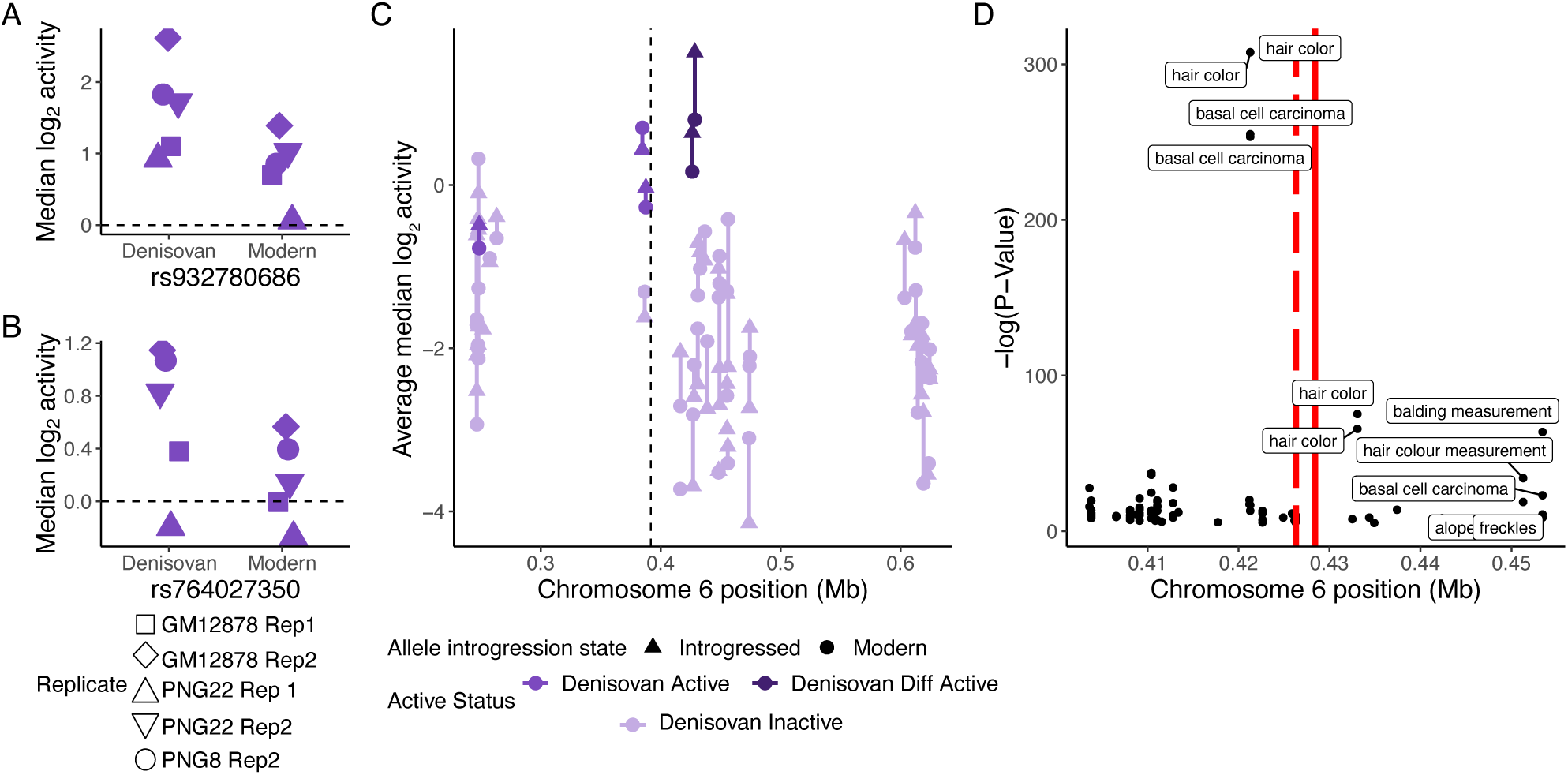
Two differentially active SNPs predicted to regulate *IFR4*. A). Activity of rs932780686 alleles across replicates. B). Activity of rs764027350 alleles across replicates. C.) Allele activity difference for SNPs surrounding rs932780686 and rs764027350. The transcription start site of *IRF4* is indicated with a dashed line. D.) GWAS catalogue hits surrounding rs932780686 (indicated with a dashed line) and rs764027350 (indicated with a solid line).

**Supplementary Figure 12.**
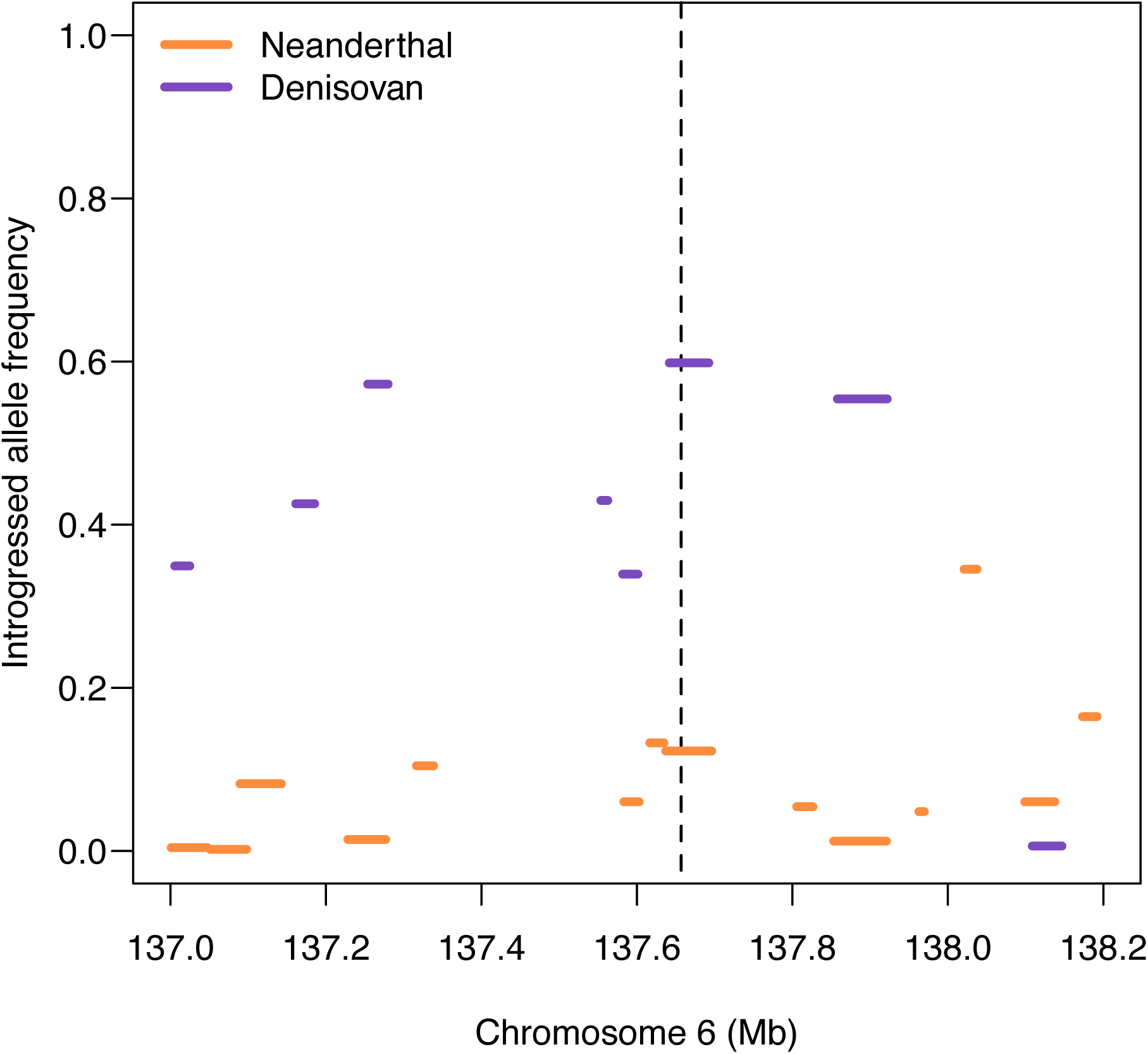
Frequency distribution of archaic haplotypes across the 1MB region surrounding rs143481175 in the Papuan dataset from Yermakovich et al., 2024. Haplotypes are classified as Neanderthal and Denisovan based on their sequence similarity with Neanderthals and the Denisovan. The position of rs143481175 is indicated by the dashed line.

**Supplementary Figure 13.**
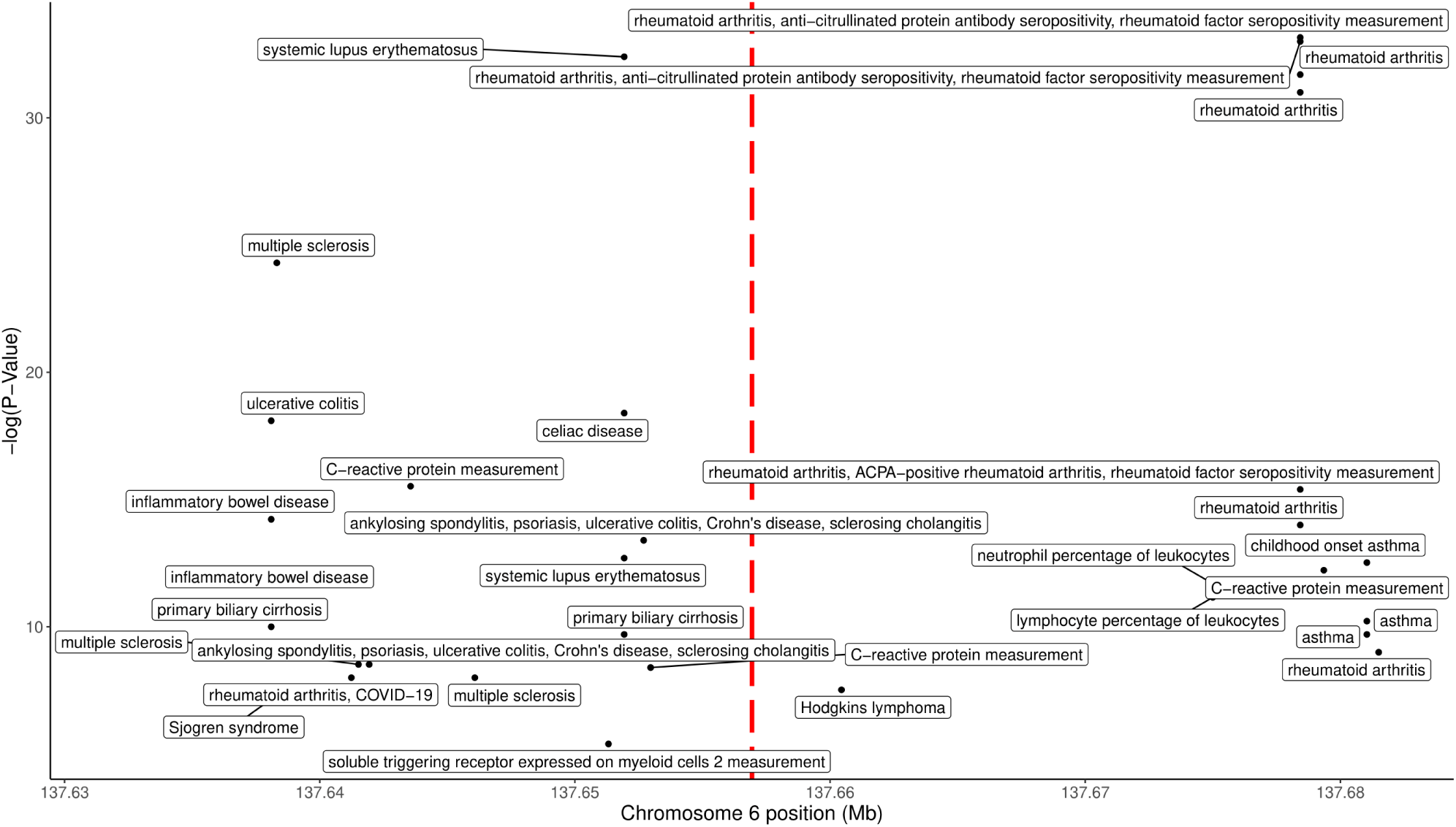
GWAS catalogue (Sollis et al., 2023) hits surrounding rs143481175. The position of rs143481175 is indicated by the red dashed line.

**Supplementary Figure 14.**
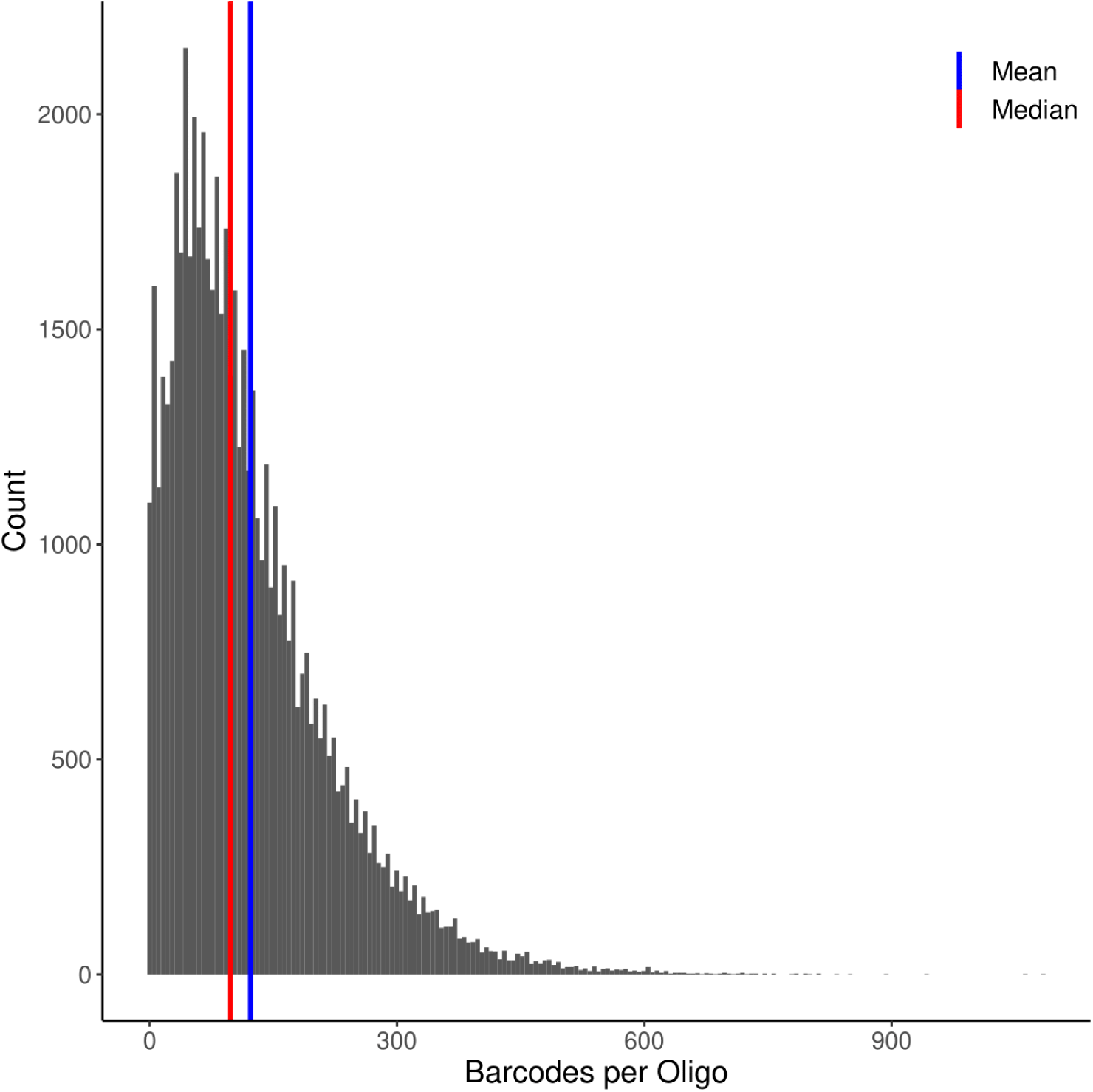
Distribution of barcodes per oligo after first round sequencing. The median number of barcodes per oligo is indicated in red, the mean per oligo in blue. Barcode counts per oligo were generated with the MPRAmatch pipeline (https://github.com/tewhey-lab/MPRA_oligo_barcode_pipeline).

**Supplementary Figure 15.**
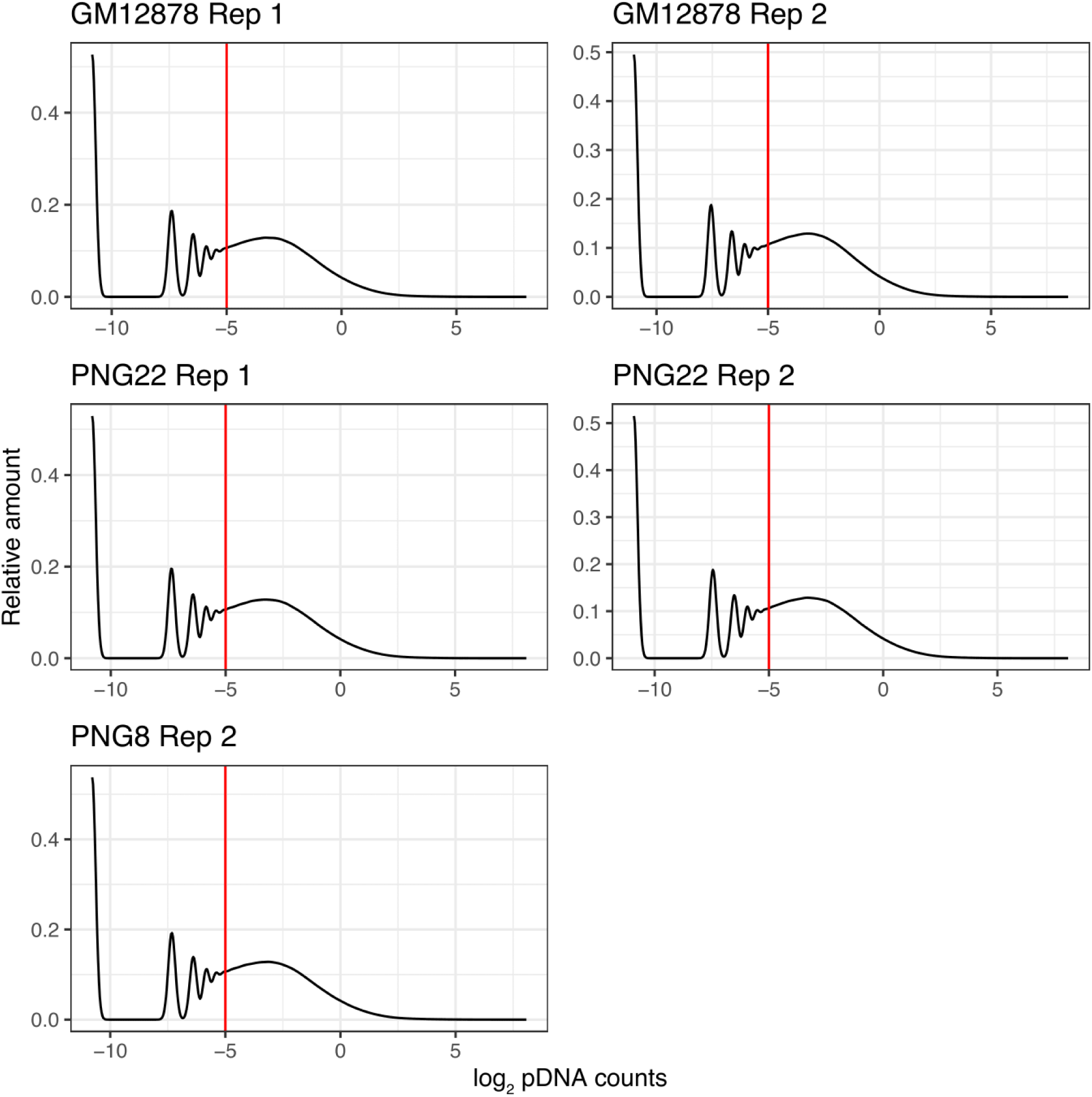
Density plots of log2 pDNA barcode counts per replicate. We excluded all barcodes with a log2CPM below -5, indicated by the red line.

## Supplementary Files

**Supplementary file 1**: All sequences included in MPRA library.

**Supplementary file 2**: Output of activity testing of single-variant sequences..

**Supplementary file 3**: Output of differential activity testing of single-variant sequences.

**Supplementary file 4**: Number of alleles per haplotype.

**Supplementary file 5**: ANOVA p-values for differential activity testing of haplotype sequences.

**Supplementary file 6**: Significant results of rGREAT enrichment testing.

**Supplementary file 7**: Primer sequences used in MPRA cloning.

**Supplementary file 8**: Output of MPRAmatch pipeline for creating barcode-oligo associations.

**Supplementary file 9**: Normalised and log2 transformed output of MPRAcount pipeline.

